# Organic dust induced mitochondrial dysfunction could be targeted via cGAS-STING or mitochondrial NOX-2 inhibition

**DOI:** 10.1101/2020.07.01.182535

**Authors:** Nyzil Massey, Denusha Shrestha, Sanjana Mahadev Bhat, Naveen Kondru, Adhithiya Charli, Locke A. Karriker, Anumantha G. Kanthasamy, Chandrashekhar Charavaryamath

## Abstract

Organic dust (OD) exposure in animal production industries poses serious respiratory and other health risks. OD consists of microbial products and particulate matter and OD exposure induced respiratory inflammation is under intense investigation. However, the effect of OD exposure on brain largely remains unknown. Recently, we have shown that OD exposure of brain microglial cells induces an inflammatory phenotype with the release of mitochondrial DNA (mt-DNA). Therefore, we tested a hypothesis that OD-exposure induced secreted mt-DNA signaling drives the inflammation. OD samples were collected from commercial swine operations and a filter sterilized OD extract (ODE) was prepared. Mouse (C57BL/6) microglial cell line was treated with medium or ODE (5%) for 48 hours along with either PBS or mitoapocynin (MA, 10 μM, NOX-2 inhibitor). Microglia treated with control or anti-STING siRNA were exposed to medium or ODE. Next, mouse (C57BL/6) pups were euthanized under an approved protocol, organotypic brain slice cultures (BSCs) were prepared and exposed to medium or ODE with or without MA treatment daily for five days. Culture supernatant, cell pellets and mt-free cytosolic fractions were processed to quantify mt-superoxide, mt-DNA, cytochrome C, TFAM, mitochondrial stress markers and mt-DNA induced signaling via cGAS-STING and TLR9. Data were analyzed using one-way ANOVA and post-hoc tests. A p value of ≤ 0.05 was considered significant. ODE exposure increased the mt-superoxide formation andMA treatment decreased the ODE-induced mt-DNA release into cytosol. ODE exposure increased the cytochrome C and TFAM levels. ODE increased MFN1/2 and PINK1 but not DRP1 and MA treatment decreased the MFN2 expression. MA treatment decreased the ODE-exposure induced mt-DNA signaling via cGAS-STING and TLR9. Anti-STING siRNA decreased the ODE-induced increase in IRF3, IFN-β and Iba1 expression. In BSCs, MA-treatment decreased the ODE induced TNF-α, IL-6 and MFN1. Taken together, OD exposure induced mt-DNA signaling could be curtailed through mitochondrial NOX-2 inhibition or STING suppression to reduce neuroinflammation.

## INTRODUCTION

Working in agriculture production and other industries is a risk factor for developing respiratory and other diseases due to exposure to contaminants (Nordgren *et al*., 2018). Among the contaminants, airborne organic dust (OD), gases (methane, ammonia, and hydrogen sulfide), odorant molecules and particulate matter (PM) of various sizes are central to the adverse health effects. OD generated in agricultural settings (swine, poultry, cattle, and other animal production units) is a complex mixture of PM and microbial products consisting of various pathogen-associated molecular patterns (PAMPs). Among the PAMPs, lipopolysaccharide (LPS) and peptidoglycan (PGN), bacterial DNA, and fungal spores have been documented in the OD samples (American Thoracic Society, 1998; Charavaryamath *et al*., 2006; Iowa State University and University of Iowa, 2002; Roy *et al*., 2003). Occupational exposure to OD is known to result in various respiratory symptoms, including bronchitis, asthma-like symptoms, coughing, sneezing, mucus membrane irritation, chest tightness, and an annual decline in lung function. Cell and molecular mechanisms underlying these respiratory exposures are under investigation by several groups (Knoell *et al*., 2019; Poole *et al*., 2019; Warren *et al*., 2019) including our laboratory (Bhat *et al*., 2019; Massey *et al*., 2019b; Nath Neerukonda *et al*., 2018).

Despite investigation of OD-exposure induced respiratory diseases, the effect of exposure to OD on other vital organs such as the brain largely remains unknown. Particularly the published observation that there is an increased incidence of Parkinson’s disease in the mid-western and north-eastern USA is interesting (Wright Willis *et al*., 2010a). These areas also know for a higher density of animal production facilities. Microglial cells of the brain are primary sentinel cells that respond to danger signals by morphological signs of activation and production of pro-inflammatory mediators (Block *et al*., 2007; Wolf *et al*., 2017). Our previous work demonstrated that OD exposure induces a pro-inflammatory phenotype in a mouse microglial cell line. Following OD exposure, microglia secreted pro-inflammatory mediators and reactive species. OD exposure also resulted in nucleocytoplasmic translocation of high-mobility group box 1 (HMGB1) and HMGB1-RAGE signaling. Following pharmacological inhibition of HMGB1 translocation using ethyl pyruvate or siRNA mediated suppression of HMGB1, we observed a reduction in reactive species, TNF-α and IL-6 production collectively. Next, pharmacological inhibition of mitochondrial NOX-2 reduced OD-induced RNS production in microglia (Massey, et al., 2019b). These results indicate a prime role for microglia in OD exposure induced neuroinflammation. Further, our results indicated that secreted HMGB1 and cellular mitochondria could be an attractive therapeutic target to curtail OD-induced neuroinflammation.

Cellular homeostasis is controlled mainly by organelles such as mitochondria and endoplasmic reticulum (ER). ER lumen maintains a unique environment which plays a vital role in protein folding. This delicate homeostasis in the lumen can be disturbed by high protein demand and inflammatory processes resulting in a large number of misfolded proteins in the ER lumen. ER stress responses are initiated by mainly by three transmembrane proteins: Inositol Requiring 1 (IRE1), PKR-like ER kinase (PERK), and Activating Transcription Factor 6 (ATF6). Activated IRE1 leads to splicing of X-box binding (XBP1) mRNA which functions as a stable UPR transcription factor. Activating transcription factor 4 (ATF4) is involved in protein folding and anti-oxidative stress (Oslowski *et al*., 2011). Glucose regulated protein 94 (GRP94), a protein that resides in the lumen of ER and regulates protein folding (Marzec *et al*., 2012).

Mitochondria are a seat of energy production and are located in the cytoplasm of cells and additionally contribute to essential cellular functions such as calcium signaling, immunity, and apoptosis. A growing number of publications indicate that mitochondrial dysfunction is central to neurodegenerative disorders (Lin *et al*., 2006) (Johri *et al*., 2012). Mitochondria are dynamic organelles that continually undergo fusion (mediated by MFN1 and MFN2 genes), fission (mediated by DRP1 gene) and several metabolic and neurological disorders are known to alter these genes (MFN1/2 and DRP1) to affect mitochondrial homeostasis (Chan, 2006) (Johri, et al., 2012). Mitochondria also undergo PINK1 mediated mitophagy during apoptosis, and mitophagy is a crucial regulator of mitochondrial turnover and cell death (Lin, et al., 2006) (Truban *et al*., 2017).

Now, growing evidence suggests a prime role for mitochondria in triggering and maintaining inflammation and mitochondrial dysfunction is emerging as a key factor in inflammatory processes (Escames *et al*., 2012). Many damage-associated molecular patterns (DAMPs) such as peptides, lipids, and mitochondrial DNA (mt-DNA) are known to be released from mitochondria and are known to act as inflammogens through activation of pattern recognition receptors (PRRs). Particularly, mt-DNA released from the mitochondrial matrix is known to signal through multiple PRRs such as Toll-like receptor 9 (TLR9), Nod-like receptors-3 (NLRP3), and cyclic GMP–AMP synthetase/stimulator of interferon gene (cGAS-STING) pathways and could mount an exaggerated inflammatory response (Nakayama *et al*., 2018). mt-DNA is a circular molecule comprising of double-stranded DNA, and human mt-DNA sequencing showed that it includes 16,569 base pairs and encodes 13 proteins. Change in mitochondrial membrane integrity increases the chances of mt-DNA release into the cellular cytosol, which can lead to an auto-immune response. (Riley *et al*., 2018) or inflammation. cGAS is a DNA sensing receptor that signals through stimulator of interferon gene (STING) and leads to the production of IFN-β. Also, STING knockdown has shown to ameliorate inflammatory responses in cultured tubular cells of the kidney following cisplatin treatment (Maekawa *et al*., 2019). cGAS-STING mediated mt-DNA signaling is an emerging target to curtail inflammation (Motwani *et al*., 2019). Apocynin (4-hydroxy-3-methoxyacetophenone) is plant-derived mitochondria targeting antioxidants. Apocynin inhibits NOX2 activity and has been studied using various *in vitro* and *in vivo* models (Jin *et al*., 2014) (Gao *et al*., 2003) (Anantharam *et al*., 2007) and been shown to be well tolerated even high doses (Anantharam, et al., 2007) (Cristóvão *et al*., 2009). Mitoapocynin (MA) has been synthesized as a more efficacious analog of apocynin by conjugating a triphenylphosphonium cation moiety via an alkyl chain with differing chain lengths (C2-C11) (Ghosh *et al*., 2016). While the neuroprotective effects of MA(C2) have been established in in-vitro models (Ghosh, et al., 2016), the long-acting MA(C11) has been more extensively used in in-vivo models.(Langley *et al*., 2017) (Dranka *et al*., 2014).

In order to understand the mechanisms of OD-induced inflammation in microglial cells, it is essential to utilize relevant models. *In vitro* models of microglial cells have been used to unravel mechanisms of neuroinflammation (Nyzil et al., 2019), and mitochondrial damage has been shown to be central in many neurodegenerative diseases (Lin, et al., 2006). Therefore, we tested a hypothesis that OD exposure of microglial cells *in vitro* and *ex vivo* organotypic brain slice culture (BSCs) induces mitochondrial and endoplasmic reticulum stress responses and resultant inflammation involves the release of mt-DNA and signaling through cGAS-STING pathway.

In the current manuscript, we demonstrated that exposure to OD induces mitochondrial and ER stress and release of mtDNA leading to cGAS-STING mediated signaling. Using anti-STING or mitochondria-targeted NOX-2 inhibitory agent MA (C2/C11), we demonstrate a reduction in OD-induced mitochondria and ER-stress and resultant inflammation.

## MATERIALS AND METHODS

### Chemicals and reagents

Dulbecco’s minimum essential medium (DMEM), fetal bovine serum (FBS), penicillin, and streptomycin (PenStrep), L-glutamine, and trypsin-EDTA were purchased from Life Technologies (Carlsbad, CA). Poly-D-Lysine (Sigma, P6407) was prepared and stored as 0.5 mg/mL stock at −20° C. Oligomycin, hydrogen peroxide, carbonyl cyanide 4-trifluoromethoxy-phenylhydrazone (FCCP) and antimycin A were purchased from Sigma Aldrich (St. Louis, MO), and the Seahorse FluxPak calibration solution was purchased from Seahorse Biosciences (Billerica, MA). Rhod-2 AM (ab142780) was purchased from Abcam (Cambridge, MA). Mitotraker green, Fluo-4AM kit, (DPBS) Dulbecco’s phosphate buffer saline, mitochondrial isolation kit (89874), DNA purification kit (Thermo fisher scientific, cat # K0512) were purchased from Thermo Fisher Scientific (Waltham, MA). IGEPAL CA630 (I3021) was purchased from Sigma. Mito-Apocynin (MA) was procured from Dr. Balaraman Kalyanaraman (Medical College of Wisconsin, Milwaukee, WI), stock solution (10 mM/L in DMSO) was prepared by shaking vigorously and stored at – 20° C. MA was used (10 µM/L) as one of the co-treatments (Table 1).

**Table 1.** Microglial Cell Treatments

| Treatment Groups | Co-treatment |
| --- | --- |
| Control | Medium |
| ODE | ODE 1% v/v |
| ODE+MA (C2/C11*) | ODE 1% v/v + MA 10 µmol |

\*C11 fraction of the MA was used to treat organotypic BSCs and C2 fraction of the MA was used in other *in vitro* experiments.

### Preparation of organic dust extract

All the experiments were conducted in accordance with an approved protocol from the Institutional Biosafety Committee (IBC protocol# 19-04) of Iowa State University. Settled swine barn dust (representing OD) was collected from various swine production units into sealed bags with a desiccant and transported on ice to the laboratory. Organic dust extract (ODE) was prepared as per a published protocol (Romberger et al., 2002). Briefly, dust samples were weighed, and for every gram of dust, 10 mL of Hank’s balanced salt solution without calcium (Gibco) was added, stirred, and allowed to stand at room temperature for 60 min. The mixture was centrifuged (1365 g, 4° C) for 20 min, supernatant collected, and the pellet was discarded. The supernatant was centrifuged again with the same conditions, the pellet discarded and recovered supernatant was filtered using a 0.22 µm filter and stored at −80° C until used. This stock was considered 100% and diluted in cell culture medium to prepare a 1% v/v solution to use in our experiments (Table 1). We routinely quantify the LPS content of our ODE samples using Pyrochrome® Kinetic Chromogenic Endotoxin (CAPE COD, Cat # CG-1500-5) as per the instructions. LPS content of several of our ODE samples has been reported in a manuscript from our group (Bhat et al., 2019).

### Cell culture and treatments

Wild-type mouse microglial cell line derived from wild-type C57BL/6 mice (Halle *et al*., 2008) was a kind gift from Dr. DT Golenbock (University of Massachusetts Medical School, Worcester, MA) to Dr. AG Kanthasamy. Microglial cells were grown in T-75 flasks (1 × 106 cells/flask), 12 well tissue culture plates coated with poly-D Lysine, 24 well tissue culture plate (50 × 10^3^ cells/well) and in 24 well seahorse assay plate (15 × 10^3^ cells/flask). Cells were maintained in DMEM supplemented with 10% heat inactivated FBS, 50 U/mL penicillin, 50 µg/mL streptomycin, and 2 mM L-glutamine. Cells were incubated overnight before treatment. All the treatment groups with pre-treatment and co-treatment details are outlined in Table 1. Control group samples were collected at 0 h because the control group samples from 6, 24, and 48 h time points did not show any differences in our pilot studies. To address the specific role of the cGAS-STING pathway, in separate experiments, cells were treated either with anti-STING or negative control siRNAs followed by either saline or ODE-exposure.

### Microglial morphology

Cells were grown in 24 well plates and maintained in Dulbecco’s Modified Eagle’s Medium (DMEM, Thermo Fisher Scientific, Walktham, MA) supplemented with 10% heat inactivated fetal bovine serum (FBS, Atlanta Biologicals, Flowery Branch, GA, cat# S11150H and lot # A17002), 50 U/mL of penicillin, 50 µg/mL of streptomycin and 2 mM L-glutamine and incubated overnight. Cells were treated as outlined in Table 1. Live cell imaging of the microglial cells was performed under an inverted bright field microscope (ALPHAPHOT-2, Nikon). Total and activated microglial cells were identified based on the morphological features (Hinwood *et al*., 2012) (Crews *et al*., 2015) (Ransohoff, 2016) and counted manually using ImageJ (NIH) in five randomly selected microscopic fields viewed under 20X objective lens. Percent of activated cells were quantified, data analyzed and graphically represented.

### Western Blot analysis

Cells were grown in T-75 flasks (1 × 10^6^ cells/flask), incubated overnight at 37° C with 5% CO_2_. Following co-treatments (outlined in Table 1), cells were treated with 0.5% trypsin for 15 min at 37° C and then re-suspended in equal volumes of DMEM and 10% FBS. Mitochondrial fraction was separated using a mitochondrial isolation kit (Thermo Fisher Scientific, catalog # 89874). Whole-cell lysates were prepared using RIPA buffer, and total protein was estimated using Bradford assay. Equal amounts of proteins (20 μg/well) were resolved on 10% SDS-PAGE gels (Bio-Rad). Next, proteins were transferred to a nitrocellulose membrane, and the nonspecific binding sites were blocked for an hour with a blocking buffer specially formulated for fluorescent western blotting (Rockland Immunochemicals, Pottstown, PA). Membranes were incubated overnight at 4 °C with the respective primary antibodies (Outlined in Table 2) namely caspase 3 Caspase 9, Cytochrome c, TFAM, MFN1, MFN2, DRP1, PINK1, SOD-2, TLR9, cGAS, IFN-β, IBA1, β-actin (1:5000, Abcam; ab6276 or ab8227). Next, membranes were incubated with the respective secondary donkey anti-rabbit IgG highly cross-adsorbed (A10043) or anti-mouse 680 Alexa Fluor antibodies (A21058, Thermo Fisher Scientific). Membranes were washed three times with PBS containing 0.05% Tween-20 and visualized on the Odyssey infrared imaging system. Using GAPDH or β-actin as a loading control, band densities were normalized, and densitometry was performed (ImageJ, NIH).

**Table 2.**
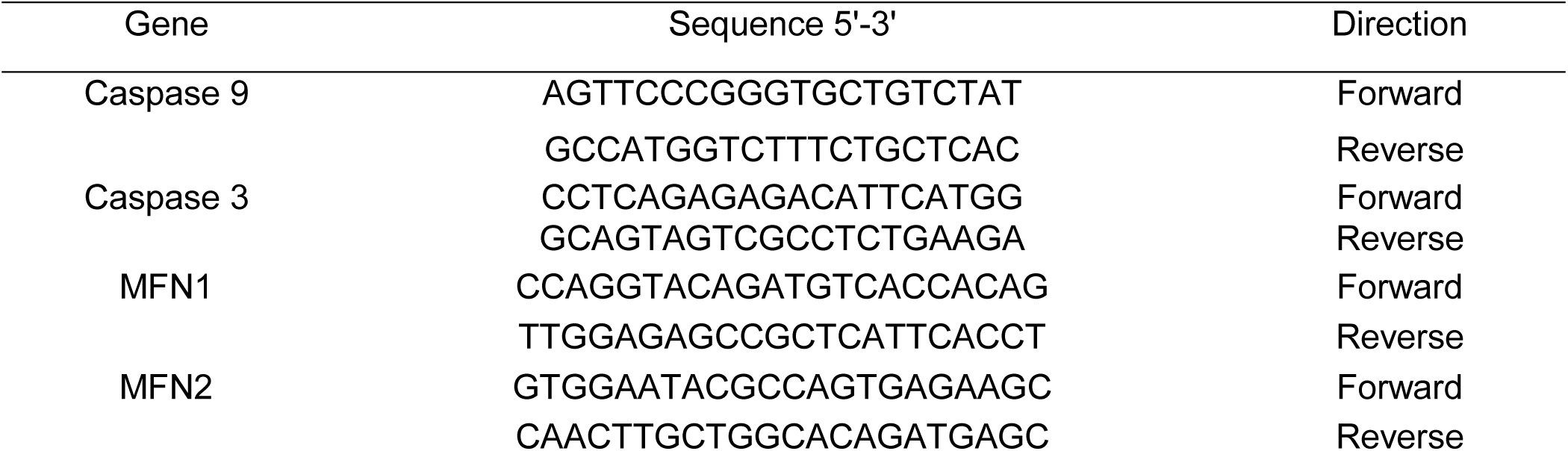
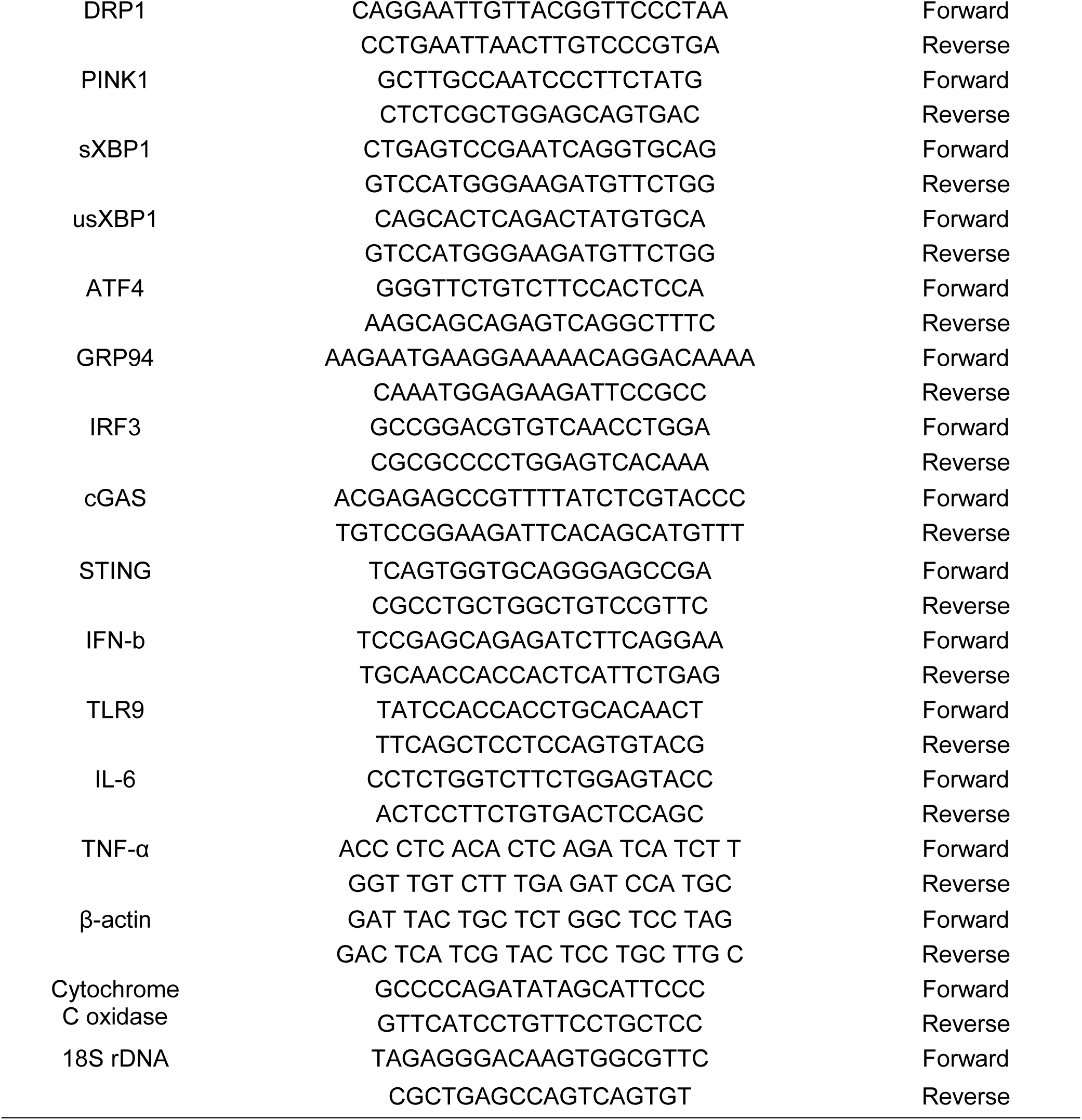
Primer Sequences

**Table 3.**
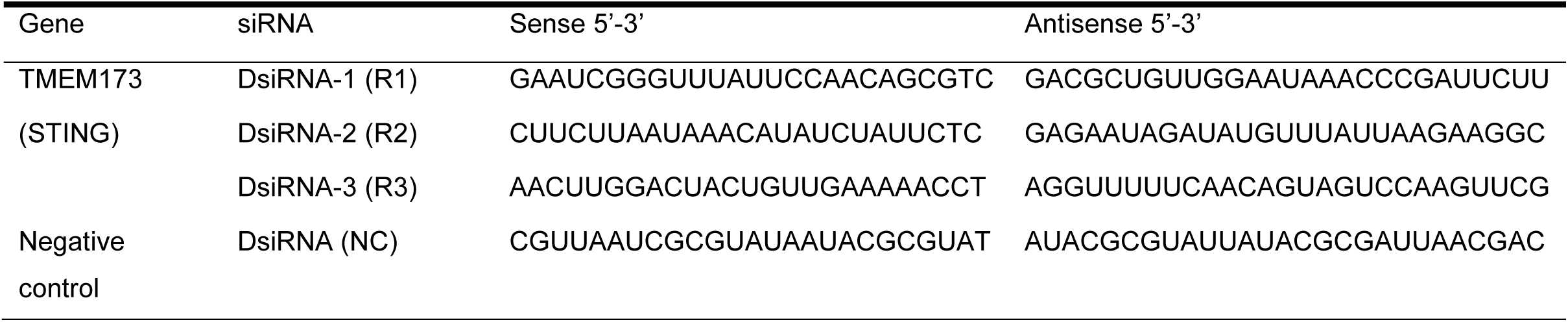
Anti-STING and Control DsiRNAs Sequences

### Transmission Electron Microscopy

Cells were grown in 12 well tissue culture plate coated with Poly-D Lysine and maintained in Dulbecco’s Modified Eagle’s Medium (DMEM, ThermoFisher Scientific, Walktham, MA) supplemented with 10% heat-inactivated fetal bovine serum (FBS), 50 U/mL of penicillin, 50 µg/mL of streptomycin and 2 mM L-glutamine and incubated overnight. Cells were treated as outlined in Table 1 and after 6, 24 and 48 h cells were washed with ice cold HBSS and fixed with 1% paraformaldehyde and 2.5% glutaraldehyde for 24 h in 4^**o**^ C. Fixed cells were embedded, sectioned, stained and imaged at the electron microscopy core facility (Roy J. Carver High Resolution Microscopy Facility, Iowa State University, Ames, IA). An investigator blinded to the treatment groups identified the ultra-structural changes in a blinded fashion.

### Seahorse assay

Mitochondrial oxygen consumption/bioenergetics was measured using a Seahorse XFe24 Extracellular Flux analyzer (Dr. Anumantha G. Kanthasamy’s laboratory, Iowa State University, Ames, IA) as described previously (Dranka *et al*., 2011). The Seahorse XFe24 Extracellular Flux analyzer is a sensitive, high-throughput instrument that performs real-time measurements of respiration rates of cells with or without oxidative stress. For quantifying mitochondrial bioenergetics, cells were first treated as outlined in Table 1 and maintained in 5% CO_2_ at 37°C for 48 h. Simultaneously, a Seahorse FluxPak cartridge was equilibrated 24 h before seahorse analysis and loaded with the mito-stressor agents like oligomycin (1 μg/mL), FCCP (1 μM) and antimycin A (10 μM). Once the mito-stressors were loaded in their corresponding position in the cartridge, the treated plate with microglia was introduced into the Seahorse analyzer covered with the FluxPak cartridge. The analyzer was then programmed to measure the basal oxygen consumption rate (OCR) readouts in five specified time intervals before progressing to inject the mito-stressors. The mito-stressors were injected after every three cycles of measurements of OCR. Further analysis of data was performed as a T-test.

### Confocal cell imaging of microglial cells

Cells were seeded into 35 mm diameter glass bottom wells and maintained in Dulbecco’s Modified Eagle’s Medium (DMEM, Thermo Fisher Scientific, Walktham, MA) supplemented with 10% heat inactivated fetal bovine serum (FBS), 50 U/mL of penicillin, 50 µg/mL of streptomycin and 2 mM L-glutamine and incubated overnight. Cells were treated as outlined in Table-1. After washing with ice cold DPBS. Cells were loaded with either mitochondria localizing probe MitoTracker^®^ Green (Invitrogen cat # M7514) (100 nM, for 12 min) followed by the fluorogenic mitochondria-targeted Ca^2+^ probe Rhod-2 AM (10 µM) for 30 min in 37°C in DMEM containing 10% fetal calf serum or MitoSox^®^ (Invitrogen cat # M36008) (100 nm) (mitochondrial superoxide staining dye) for 15 min. After loading with either of the dyes, cells were placed on the stage of the Nikon microscope. MitoTracker^®^ Green was observed using FITC filter. Rhod-2 AM and MitoSox^®^ were observed using cy3 filter. Following MitoSox staining, five fields (under a 20X objective lens) per slide were chosen randomly, and total stained cells were counted (ImageJ, NIH). Next, the total staining intensity per field (cy3) was measured using software (HC Image, Hamamatsu Corp, Sewickley, Pennsylvania). The total intensity of the microscopic field (cy3) was divided by the total number of cells per field to obtain mean intensity (cy3) per cell.

### qRT-PCR

Total cellular RNA was isolated using TRIzol™ (Invitrogen cat # 15596-026) extraction methods as per manufacturer’s guidelines. Following treatment (Table 1), RNA concentration was measured using NanoDrop, and the A260/A280 ratio assessed quality. Samples with an A260/A280 ratio between 1.8-2.1 were considered acceptable and used for further analysis. One microgram of RNA was reverse transcribed into cDNA using the superscript IV VILO Kit (Thermo Fisher Scientific, cat # 11766050) following the manufacturer’s protocol. For qPCR, 5 µL of PowerUp™ SYBR Green Master mix (Thermo Fisher Scientific, cat # 25742), 0.5 µL each of forward and reverse primers (10 µM), 3 µL of water and 1 µL of cDNA (1-10 ng) were used. The genes and their respective primer sequences used for qRT-PCR analysis are listed in Table 2, and all the primers were synthesized at Iowa State University’s DNA Facility. 18 S and β-actin were used as a housekeeping gene in all the qRT-PCR experiments. No-template controls and dissociation curves were run for all experiments to exclude cross-contamination. *C*_T_ values of gene products of interest were normalized to housekeeping gene product *C*_T_ values. Comparisons were made between experimental groups using the Δ*C*_T_ method. Briefly, the Δ*C*_T_ value was calculated for each sample (*C*_T_ gene of interest minus *C*_T_ 18 S or β-actin). Then the calibrator value was averaged (Δ*C*_T_) for the control samples. The calibrator was subtracted from the Δ*C*_T_ for each control and from the experimental sample to derive the ΔΔ*C*_T_. The fold change was calculated as 2^−ΔΔCt^. Average fold change was calculated for each experimental group.

### Mitochondrial DNA detection

Cells were seeded in T-25 tissue culture flask (2x 10^5^ cells/flask). After administering the treatments, cells were washed once with ice-cold DPBS and lysed with 1% Nonidet P-40 (Igepal Ca-630, Sigma-Aldrich, cat#18896). Following centrifugation (13000 rpm for 15 min at 4° C), mitochondria free cytosolic fraction of microglia was collected, and DNA was purified using DNA purification kit (Thermo fisher scientific, cat # K0512). For q-PCR, 10 µL of SYBR Green Mastermix (Qiagen Cat #208056), 1 µL of primers, 8 µL of water and 1 µL of cDNA were used. The cytochrome c oxidase gene was used in our qRT-PCR reaction (primer sequences: forward 5’-GCCCCAGATATAGCATTCCC-3, and reverse 5’-GTTCATCCTGTTCCTGCTCC-3’, synthesized at Iowa State University’s DNA Facility) (Bronner *et al*., 2016) to quantify the amount of mt-DNA. Using primers specific for 18S, we quantified the housekeeping gene expression.

### siRNA-mediated knockdown of STING

Our protocol for siRNA mediated knockdown of a gene of interest is published (Massey, et al., 2019b). Briefly, three (R1, R2, and R3) custom-designed Dicer-substrate anti-STING siRNAs (DsiRNA), scrambled RNA (negative control, NC), HPRT1 DsiRNA (positive control), and a fluorescent transfection control (TYE563) were purchased from Integrated DNA Technologies Inc. (Coralville, Iowa) to maximize the probability of achieving successful STING knockdown. Lipofectamine 2000 (Thermo Fisher Scientific) was employed to transfect DsiRNA into the microglia. Sequences of siRNAs and NC siRNA are listed in Table-4. For performing transfection with various siRNAs, microglia were cultured a day before in DMEM without antibiotics and FBS. For each transfection, 20 mmol of (DsiRNA) for STING, scrambled RNA (negative control), HPRT1 DsiRNA (positive control), and a fluorescent transfection control (TYE563) were diluted in Opti-MEM media without antibiotics and FBS to 10 nmol and gently mixed with Lipofectamine 2000 according to the manufacturer’s protocol. Following incubation for 20 min at room temperature, the transfection mixture was added to the cells, transfected cells were further incubated at 37 **°**C for 24, and 48 h, and knockdown was confirmed by qRT-PCR and Western blot analysis of the target gene and protein respectively. Transfection was confirmed by performing immunofluorescence (ICC) for TYE563 (fluorescent transfection control).

### DNase treatment of ODE

In order to eliminate any interference by the prokaryotic or eukaryotic DNA that may be present in 0.2 µm filtered ODE samples, we performed the DNase treatment. Total RNA in an ODE sample was measured using NanoDrop (Biotek synergy 2). Then RNA concentration was used as a criterion for the amount of DNase (Turbo Dnase, Thermo Fisher Scientific cat # AM2239) to be added in 50µl volume of 100% ODE to remove any traces of DNA effectively.

### Mito-Apo treatment of microglia and brain slice cultures (BSCs)

We did not use mito-apocynin treatment whenever there was no ODE exposure effect. Two forms of Mito-apocynin (C2 or C11) were used in this study. Unless otherwise specified, MA refers to C2 fraction. Details of the preparation of MA (C2 and C11) and its neuroprotective properties have been reviewed (Ghosh, et al., 2016) (Dranka, et al., 2014). For treatment purposes, in-vitro model wild type microglial cell model was treated with MA i.e., C2 form of MA at 10 µmol for 48 h (Ghosh, et al., 2016). Since, MA(C11) is the preferred form for in-vivo model (Dranka, et al., 2014). MA(C11) was also found to be more neuroprotective on ex-vivo model at 10 µmol for 5 day as compared to MA(C2) (data not shown). MA (C2 or C11) exposure for In-vitro and ex-vivo models was always done as co-treatments with ODE (1%). MA(C2 or C11) alone in media was never done as apocynin has no toxic effect on cell culture and mouse models and can be well tolerated even at higher doses (Anantharam, et al., 2007) (Cristóvão, et al., 2009).

### Brain slice cultures (BSCs)

All the work described here was performed as per the approved protocols from the Iowa State University’s Institutional Animal Care and Use Committee (IACUC protocol # 18-290). Organotypic slices were prepared as previously described (Kondru *et al*., 2017). As per an approved protocol (IACUC protocol#18-227), we procured the breeding pairs of C57BL/6 mice (The Jackson Laboratories, Bar Harbor, ME) and paired male and female mice of around four weeks of age. Mouse pups born were kept with the parents until 9-12 days. Mouse pups (WT, C57BL/6, 9-12 days, male or females) were euthanized by cervical dislocation by pinching and disrupting in the high cervical region as per an approved AVMA method for the euthanasia of pups of 9-12 days of age. Organotypic brain slices (BSCs) were prepared from freshly dissected brain tissues using a microtome (Compresstome(tm) VF-300, Precisionary Instruments). After dissection, the whole brain was oriented in the mid-sagittal plane in the Compresstome’s specimen tube, which had been prefilled with 2% low-melting-point agarose. The agarose was quickly solidified by clasping the specimen tube with a chilling block. Then, the specimen tube was inserted into the slicing reservoir filled with freshly prepared, ice-cold Gey’s balanced salt solution supplemented with the excitotoxic antagonist, kynurenic acid (GBSSK). To prepare GBSS, we added the following in solution in the following order from 10x stocks to obtain the final concentrations per liter: 8 g NaCl, 0.37 g KCl, 0.12 g Na_2_HPO_4_, 0.22 g CaCl_2_ · 2H_2_O, 0.09 g KH_2_PO_4_, 0.07 g MgSO_4_ · 7H_2_O, 0.210 g MgCl_2_ · 6H_2_O, 0.227 g NaHCO_3_. The compression lip located in the cutting chamber helps to stabilize the brain specimen while obtaining 350-μm thick slices with the blade set at a medium vibration speed. BSCs were collected at the specimen tube’s outlet and transferred to another plate with fresh prefilled GBSSK. Later, the BSCs were washed twice in 6 mL ice-cold GBSSK, transferred to 6-well plate inserts (Falcon #353090, 3-4 slices per insert), and were incubated in a humidified atmosphere with 5% CO_2_ at 37 °C. Culture media was replenished every other day for two weeks before starting any treatments. After two weeks of incubation, BSCs cultures were treated for five days. All the treatment groups with pre-treatment and co-treatment details are outlined in Table 1.

### Confocal imaging of BSCs

After five days of treatment with either control or ODE followed by either MA (C11) or vehicle (PBS) BSCs on inserts were cut out with a scalpel and placed in a new 12 well inserts facing upward and washed twice with phosphate-buffered saline (PBS) and fixed in 4% paraformaldehyde at room temperature for 30 min and incubated with ice-cold 20% methanol in PBS for an additional 5 min. BSCs were permeabilized with 1% Triton X-100 in PBS for 12-18 h at 4°C. Blocking was performed with 20% BSA with 0.1 % Triton X-100 in PBS for 2-3 h. Next, BSCs were incubated with anti IBA-1 and anti NeuN antibodies (listed in Table 5) overnight at 4°C. After being washed with washing solution (5% BSA in PBS), BSCs were incubated with secondary antibodies (Table 4), 12 h at 4° C followed by mounting with VECTASHIELD antifade mounting medium containing 4’, 6-Diamidino-2-Phenylindole, Dihydrochloride (DAPI, Vector Labs, Burlingame, California) and covered with a cover-glass. Nuclei were stained blue with DAPI. The cover glasses were sealed with nail polish, and slides were imaged under Nikon Eclipse TE2000-U and photographed using Photometrics Cool Snap cf, HCImage (Tucson, AZ).

**Table 4.** Antibodies used in Western blotting

| Antibody | Catalogue | Dilution | Vendor |
| --- | --- | --- | --- |
| SOD-2 | Sc-133134 |  | Santa Cruz biotechnology |
| Cytochrome c | Sc-13156 |  | Santa Cruz biotechnology |
| TFAM | Sc-166965 |  | Santa Cruz biotechnology |
| TLR9 | Sc-47723 |  | Santa Cruz biotechnology |
| cGAS | Sc-515802 |  | Santa Cruz biotechnology |
| IFN-β | Sc-57201 | | Santa Cruz biotechnology |
| IBA-1 | Ab-5076 |  | Abcam |

**Table 5.** Antibodies used in ICC

| Primary Target | Secondary Antibody | Source (Catalogue Number) |
| --- | --- | --- |
| Iba1 (Goat Pab)* | Anti-Goat (FITC conjugated) raised donkey | Jackson ImmunoResearch (705-095-147) |
| NueN (Mouse Pab) | Anti-Mouse (Cy3 conjugated) raised in donkey | Jackson ImmunoResearch (711-165-150) |

\* Pab: polyclonal antibody

### TUNEL assay

After five days of treatment with either control or ODE followed by either MA (C11) or vehicle (PBS), BSCs on inserts were cut out with a scalpel and placed in a new 12 well inserts facing upward (BSCs side up) and washed twice with phosphate-buffered saline (PBS) and fixed in 4% paraformaldehyde at room temperature for 30 min. TUNEL assay was performed using a DeadEnd(tm) Fluorometric TUNEL System according to the manufacturer’s instructions (Promega Corporation, Madison, WI, USA). Briefly, BSCs were rinsed in PBS and incubated in 20 *μ*g/mL proteinase K for 10 min. After rinsing in PBS (0.05 M phosphate buffer containing 0.145 M sodium chloride, pH 7.4), the BSCs were incubated with equilibration buffer and then TdT enzyme in a humidified chamber at 37°C for 60 min. Next, BSCs were transferred into pre-warmed working strength stop wash buffer for 15 min. Following rinsing with PBS, the BSCs were mounted with brain slices facing up from with VECTASHIELD antifade mounting medium containing 4’, 6-Diamidino-2-Phenylindole, Dihydrochloride (DAPI, Vector Labs, Burlingame, California) and covered with a cover-glass. The cover glass was sealed with nail polish. Nuclei were stained blue with DAPI and localized green fluorescence of apoptotic cells was detected by fluorescence microscopy and photographed (Nikon Eclipse TE2000-U, Photometrics Cool Snap cf, HCImage). TUNEL positive cells were counted manually using ImageJ (NIH) in five randomly selected microscopic fields viewed under 20X objective lens. Percent of TUNEL positive cells were quantified, statistically analyzed, and plotted.

### Statistical analysis

Data were expressed as mean ± SEM and analyzed by T-test or one-way or two-way ANOVA followed by Bonferroni’s post hoc comparison tests (GraphPad Prism 5.0, La Jolla, California). A p-value of < .05 was considered statistically significant. An asterisk (*) indicates a significant difference between controls and ODE treated cells whereas hashtag (#) indicates either MA(C2/C11) treatment or siRNA-mediated STING knockdown effect. The p-values (Figs. 1–12 and legends) corresponding to asterisk/s or hashtag/s are listed in Table 6.

**Table 6.** Symbols (Asterisk or Hashtag) and Corresponding p-Values

| Symbol | p-value |
| --- | --- |
| * or # | $\leq .05$ |
| ** or ## | $\leq .01$ |
| *** or ### | $\leq .001$ |
| **** or #### | $\leq .0001$ |

**Fig 1.**
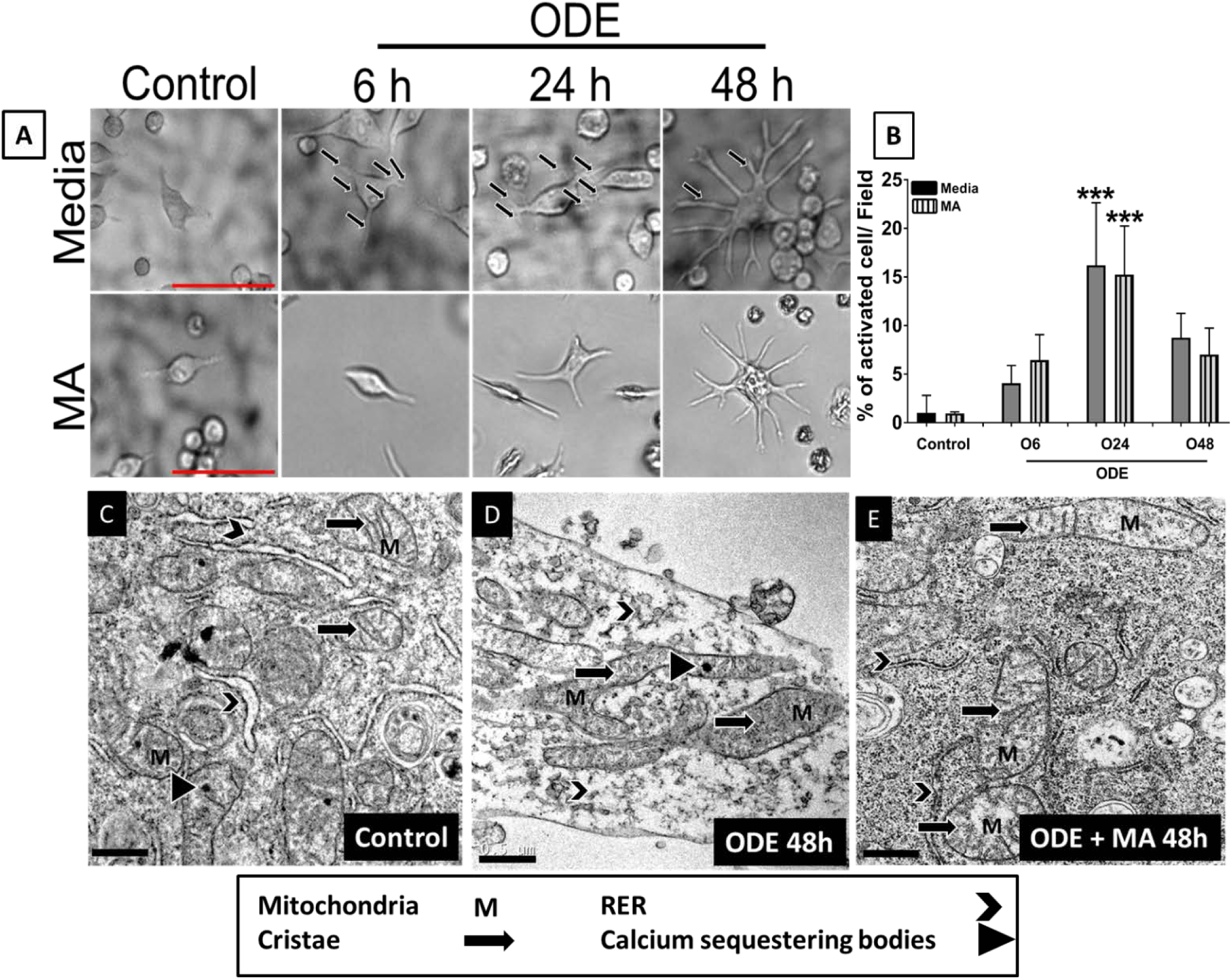
MA reduces ODE induced morphological and ultrastructural changes in Microglia. Microglia were exposed to media or ODE and co-treated with either vehicle or MA. Morphological signs of activation such as an increase in size and change in shape (amoeboid body with thick and longer processes) were observed, and percent of activated microglia/microscopic field was calculated manually at 6, 24, and 48 h. Compared to controls, ODE treated microglia appeared to increase in size with a round center and thick amoeboid processes (arrows) (A, upper panel). MA (mitochondria targeting NOX-2 inhibitor) treatment did not reduce ODE-induced changes in microglia (micrometer bar= 50 µm) (A, lower panel). The number of microglia was manually counted in five randomly chosen fields, and percent activated cells was calculated. Compared to controls, ODE significantly induced morphological signs of activation at 24 and 48 h. MA treatment had no effect on ODE-induced microglial activation at 24 and 48 h (B). Microglia were exposed to media or ODE with or without MA and processed for transmission electron microscopy (TEM) (B). Control cells showed mitochondria with normal cristae, electron-dense calcium sequestration bodies, and rough endoplasmic reticulum (RER) (C). Following ODE exposure, mitochondria were hypertrophic with cristolysis, contained larger calcium sequestration bodies, and fragmented RER (D). Compared to vehicle treatment, MA-treatment reduced ODE-induced mitochondrial hypertrophy, cristolysis, larger electron-dense calcium sequestration bodies, and fragmentation of RER at 48 h (E) (micrometer bar = 0.5 µm). (n=4, * exposure effect, P < 0.05).

## RESULTS

### ODE exposure of microglia induces microscopic and ultrastructural changes indicating mitochondrial and ER damage

Using bright field and transmission electron microscopy (TEM), we assessed the microscopic and ultrastructural changes in microglia following exposure to ODE with or without MA treatment. Compared to controls, ODE exposure of microglia resulted in morphological signs of activation as indicated by an increase in size over time (6, 24, and 48 h), and microglia developed an amoeboid body with thick and longer processes (Fig. 1A). Later, manually analyzing the microglia under a microscope indicated an increase in the number of activated microglia upon ODE treatment as compared to control at 24 h. MA did not show any effect on morphological signs of microglial activation (Fig. 1B). We also assessed the ultrastructural details of microglia using TEM. ODE treated microglia showed mitochondrial hypertrophy with cristolysis, increased calcium sequestering bodies, and fragmented RER as compared to microglia treated with vehicle. MA treatment partially reduced ODE induced changes inside microglia (Fig. 1C-E).

### ODE impairs mitochondrial function

We assessed mitochondrial bioenergetics in microglia exposed to media alone or ODE with or without MA treatment using sea horse assay. ODE treatment significantly affected the mitochondrial bioenergetics by decreasing mean OCR, basal respiration, and ATP production. MA only partially recuperated the mitochondrial bioenergetics by increasing basal respiration and ATP production (Fig. 2 A-E).

**Fig 2.**
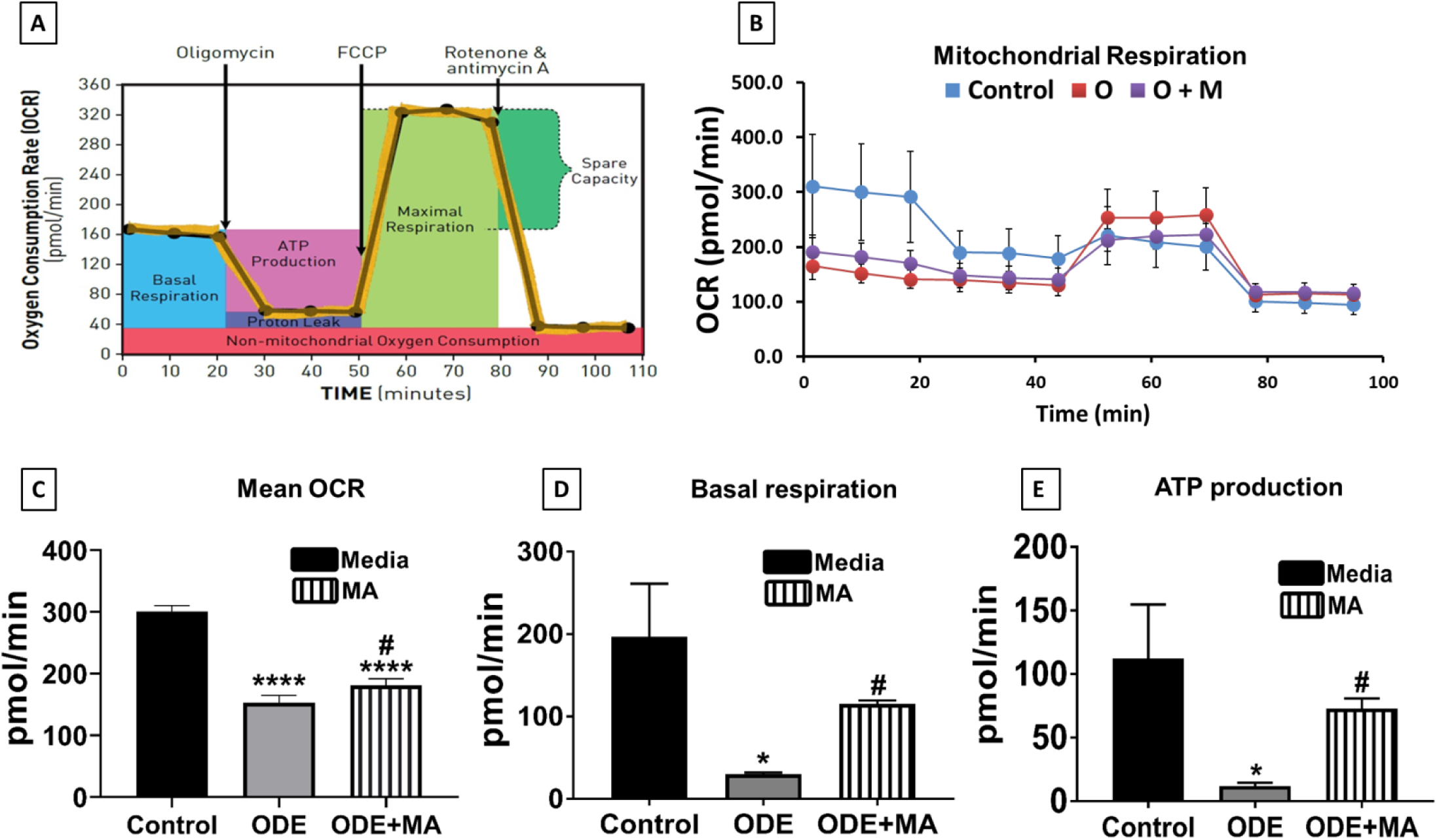
MA reduces ODE induced mitochondrial dysfunction. Cells were exposed to media and ODE with or without MA and processed for seahorse assay to measure the mitochondrial bioenergetics. Standard mitochondrial stressors (oligomycin 1 µg/mL, FCCP 1 µmol, and antimycin A 10 µmol) were used, and mitochondrial bioenergetics was measured (A). Time-lapse visualization of change in mitochondrial respiration of microglia exposed to media or ODE and co-treated with either vehicle or MA upon treatment with mitochondrial stressors (B). Compared to control, ODE treated cells showed a decrease in mean oxygen consumption rate (OCR), basal respiration, and ATP production in mitochondria. MA treatment significantly increased the mean OCR, basal respiration, and ATP production of mitochondria (C-E). (n=4, * exposure effect, # MA treatment effect, P < 0.05).

### ODE upregulates mitochondrial and endoplasmic reticulum (ER) stress responses in activated microglia

Mitochondrial and ER phenotypic changes in microglia were analyzed in microglia after exposure to ODE with or without MA treatment using qRT-PCR analysis. MFN1 mRNA levels increased by 2 folds after 48 h of ODE treatment, whereas MA had no effect on MFN1 gene activity (Fig. 3 A). MFN2 activity was also upregulated by 3 folds upon ODE treatment again at 48 h as compared to control. In contrast, MA significantly declined the mRNA levels after ODE exposure to the control levels (Fig. 3 B). On the other hand, we did not see any significant changes in the levels of DRP1 mRNA with ODE treatment with or without MA (Fig. 3 C). PINK1 gene expression significantly increased with a significant fold increase in PINK1 mRNA levels at 48 h post ODE exposure. MA treatment had no significant effect on the PINK1 gene activity (Fig. 3 D). Upon ODE treatment spliced XBP1 mRNA were found to be significantly increased as compared to control at 48 h, whereas Un-spliced XBP1 mRNA remains unaffected. MA significantly reduced spliced XBP1 mRNA levels (Fig. 3 E-F). ATF4 mRNA levels also saw a significant fold change increase upon ODE treatment 48 h post ODE treatment, whereas MA showed no effect on ATF4 levels (Fig. 3 G). The upregulation of GRP94 reflects endogenous UPR activation. GRP94 mRNA levels were significantly increased upon ODE treatment at 6h, and MA showed no effect on GRP94 mRNA levels (Fig. 3 H).

**Fig 3.**
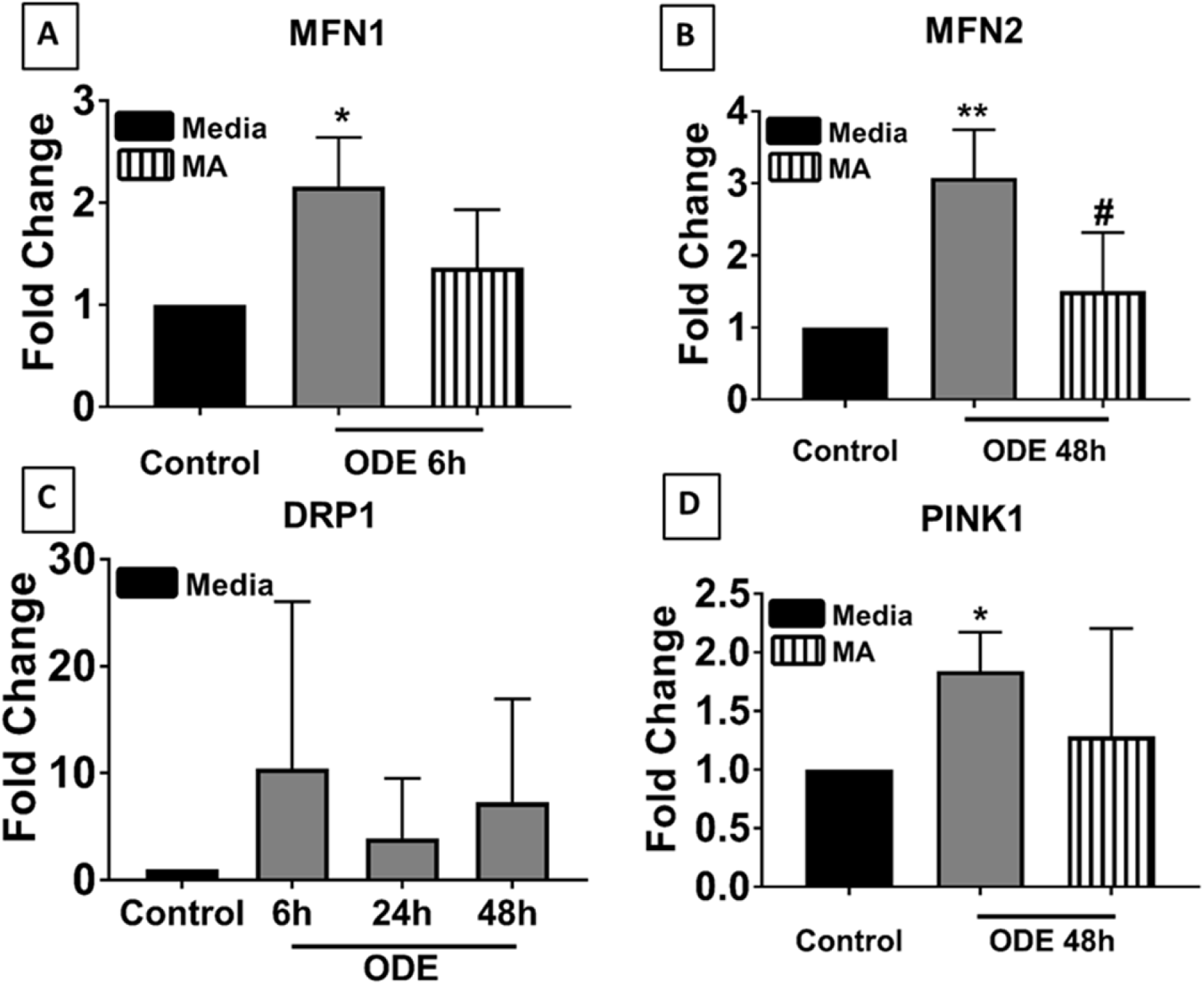
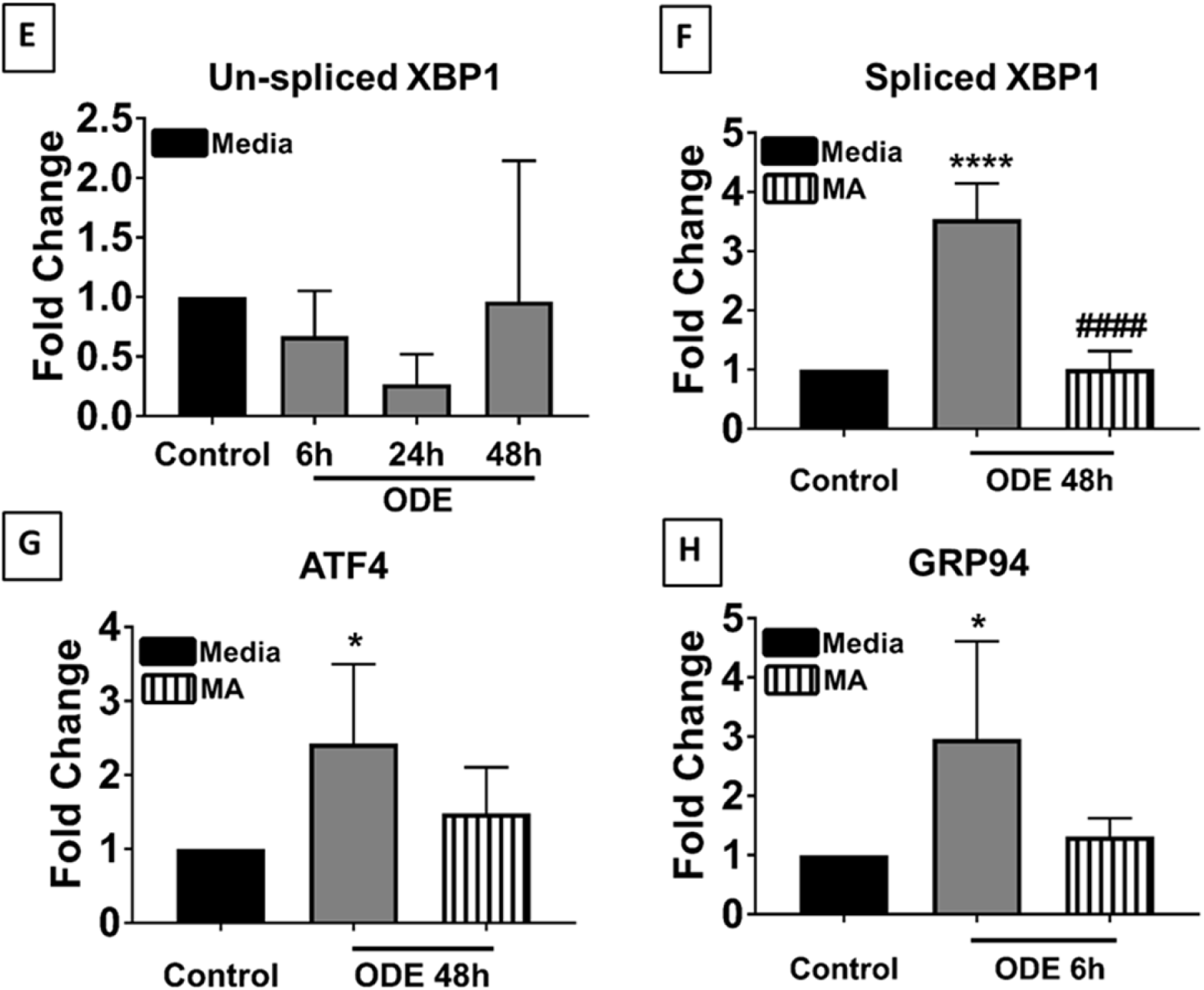
ODE induces increased expression of mitochondrial and endoplasmic reticulum stress genes. Cells were exposed to media or ODE and co-treated with either vehicle or MA and processed for qRT-PCR analysis. Compared to control, ODE exposed cells showed increased expression of MFN1 at 6 h, and MA treatment had no effect (A). Compared to control, ODE exposed cells showed increased expression of MFN2 at 6 h (data not shown), 24 h (data not shown), and 48 h following ODE treatment. MA significantly reduced MFN2 expression at 6h (data not shown) and 48 h (B). DRP1 expression did not change between control and any of the treatment groups (C). Compared to control, ODE exposed cells showed increased expression of PINK1 at 48 h, and MA treatment had no effect (D). Un-spliced XBP1 expression did not change between control and any of the treatment groups (E). Compared to control, ODE exposed cells showed increased expression of spliced XBP1 gene at 24 (data not shown) and 48 h. MA significantly reduced the expression of spliced XB1 at 48 h (F). Compared to controls, ODE exposed cells showed increased ATF4 gene expression at 6 h (data not shown) and 48 h. MA treatment did not affect ATF4 expression (G). Compared to controls, ODE exposed cells showed increased GRP94 gene expression at 6 h. MA treatment did not affect (H). (n=3 in duplicates * exposure effect, # MA treatment effect, P < 0.05).

### ODE induces reactive oxygen species (mt-ROS) production and calcium uptake by mitochondria

Fluorescent staining of microglia with mitosox® (representing mt-ROS) was done after treatment with media alone or ODE with or without MA treatment. Compared to control, ODE treatment triggered the mt-ROS production in microglia, as indicated by increased mitosox® intensity (24 h and 48 h), and MA treatment did not have any effect on the production of mt-ROS (Fig. 4 A). Also, quantification of 5 random fields (images not shown) revealed that staining intensity per cell was higher in ODE treated (24 h and 48 h) microglia as compared to control, and MA had no significant effect (Fig. 4 B). Expression of SOD-2 was measured using western blot analysis. ODE exposure increased the SOD-2 protein levels inside microglia as compared to control (24 h and 48h), MA had no significant effect on SOD-2 protein expression (Fig. 4 C-D). Colocalization of rhodamine (calcium indicator) and mitotraker® green (mitochondria marker) in ODE treated microglia compared to control at 24 h and 48 h was suggestive of increased calcium levels inside mitochondria (Fig. 4 E).

**Fig 4.**
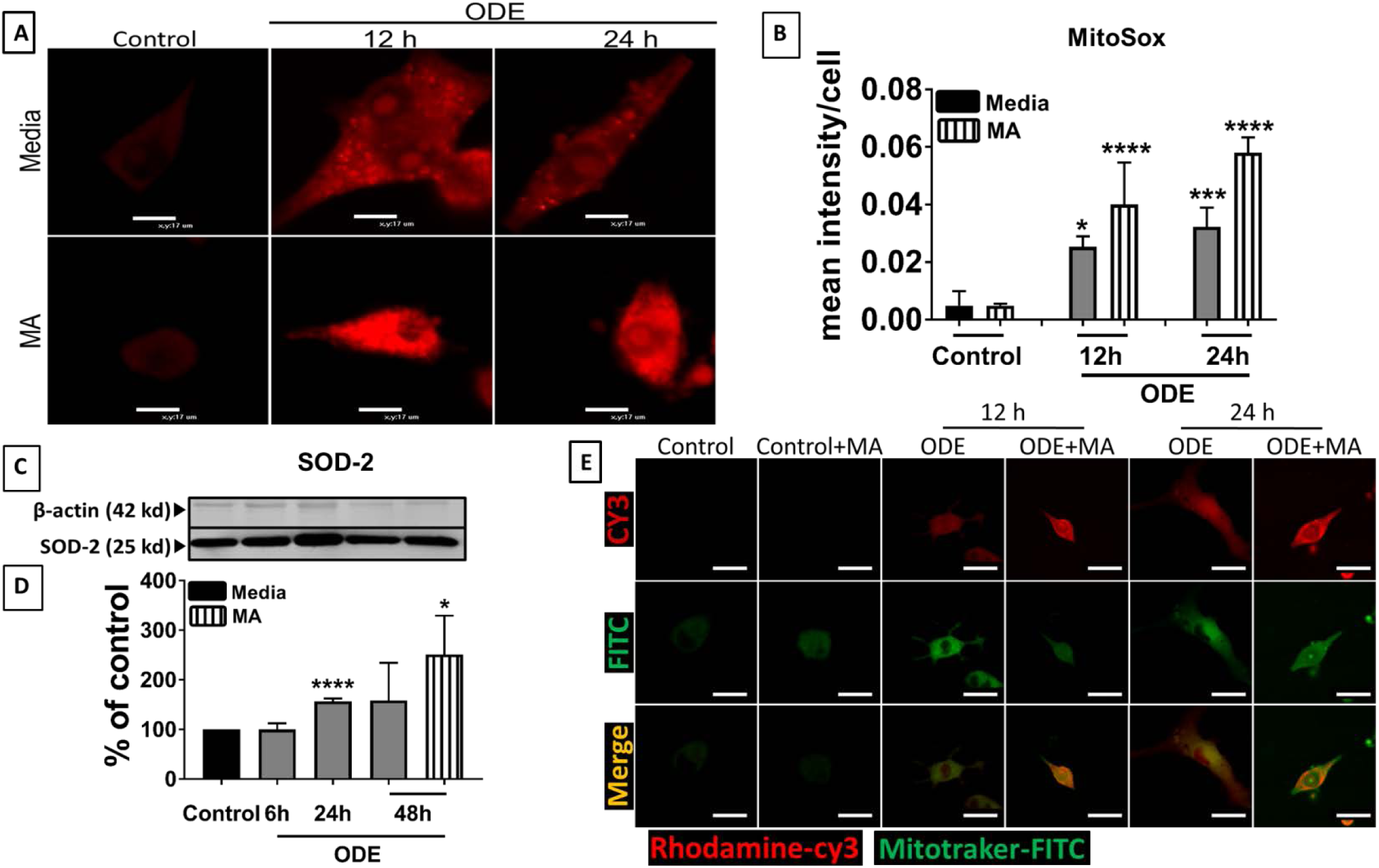
ODE induces mitochondrial superoxide production and increased mitochondrial calcium uptake in Microglia. Microglia were exposed to media or ODE and co-treated with either vehicle or MA and were stained with MitoSox® (mitochondrial superoxide) (A-B), Rhodamine (calcium marker) (E) or processed for western blot analysis using whole cell lysates (C-D). Compared to controls, ODE exposed cells appeared to contain higher amounts of MitoSox® stain (A, upper panel). Compared to the vehicle, MA treatment did not change MitoSox® staining (A, lower panel) (micrometer bar= 17 µm). Mean staining intensity in five randomly chosen fields was measured (images not shown) using computer software (HCImage, HM), and mean staining intensity per cell was calculated and plotted. Compared to controls, ODE significantly increased MitoSox® staining intensity at 24, and 48 h and MA treatment did not affect (B). Bands were normalized (GAPDH, loading control), densitometry was performed (ImageJ, NIH), and statistical analysis was conducted (n=4). Compared to controls, ODE treatment significantly increased the superoxide dismutase-2 (SOD-2) protein expression at 24 h. MA treatment elevated the SOD-2 levels at 48 h (C-D). Compared to controls, ODE exposed cells showed an increase in rhodamine dye (CY3, red-colored cytosolic calcium marker, Upper panel) and MitoTracker (FITC, green, mitochondrial marker, Middle panel) at 12 and 24 h (merged lower panel) (micrometer bar= 50 µm). MA treatment had no effect on mitochondrial calcium uptake (E) (n=3 *exposure effect, # MA treatment effect, P < 0.05).

### ODE exposure of microglia upregulates apoptotic pathway through cytochrome c release in cytosol

Western blot analysis was done for analyzing the protein expression of cytochrome c in cytosol. Mitochondria free cytosol of microglia was prepared and cytochrome c protein levels were found to be elevated upon ODE treatment compared to control at 48 h. MA treatment did not significantly reduce Cytochrome c levels after ODE treatment (Fig. 5 A-B). Furthermore, qRT-PCR analysis was performed for Caspase 3 and Caspase 9 activity. The gene expression of Caspase 3 and caspase 9 was found to be significantly elevated reflected by their mRNA levels in ODE treated microglia, and this time MA treatment successfully downregulated Caspase 3 and Caspase 9 gene activity (Fig. 5 C-D).

**Fig 5.**
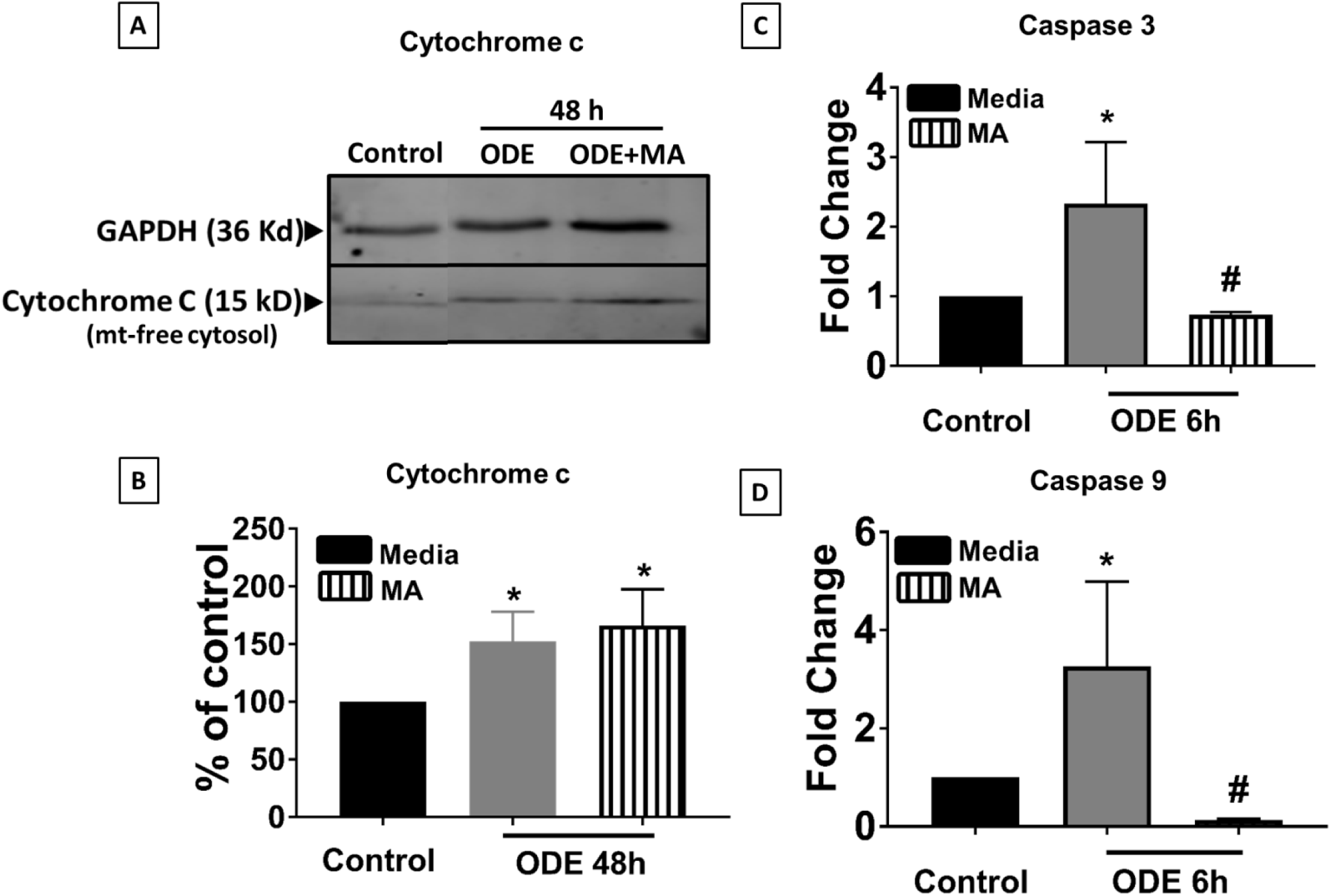
ODE treatment induces the release of Cytochrome c into the cytosol which involves Caspase 3 and Caspase 9 activation. Cells were exposed to media or ODE and co-treated with either vehicle or MA and processed either for western blot or qRT-PCR analysis. Western blot analysis of cells exposed to medium or ODE followed by vehicle or MA was performed (A-B). Normalized (GAPDH, loading control) bands of cytochrome c were processed for densitometry (ImageJ, NIH), and statistical analysis was performed. Compared to controls, ODE exposure significantly increased cytochrome c levels at 48 h and MA treatment had no effect. Compared to controls, ODE exposed cells showed an increase in the expression of caspase 3 (C) and caspase 9 (D) genes at 6 h and MA treatment had no effect on either the caspase-3 or -9 gene expression levels (C-D). (n=4 and duplicates for qRT-PCR, * exposure effect, # MA treatment effect, and p < 0.05).

### ODE induces mt-DNA and TFAM release into cytosol

Mitochondria free cytosol were prepared of microglia exposed to ODE with or without MA treatment. Upon western blot analysis, TFAM was found in the cytoplasm of microglia after 48 h post-treatment, and MA treatment could not rescue the release of TFAM from mitochondria (Fig. 6 A-B). Next, to rule out ODE as the external source of mt-DNA in our sample. We ran a qRT-PCR on ODE either treated with DNase (negative control) or without DNase. Compared to the cytosol of control microglia, ODE contained significantly fewer levels of mt-DNA (Fig. 6 C). Finally, the mt-DNA levels in the cytosol of ODE treated microglia were analyzed and found to be significantly elevated at 12 h post-treatment as compared to control. Interestingly, MA prevented the release of mt-DNA into the cytosol, unlike TFAM (Fig. 6 D).

**Fig 6.**
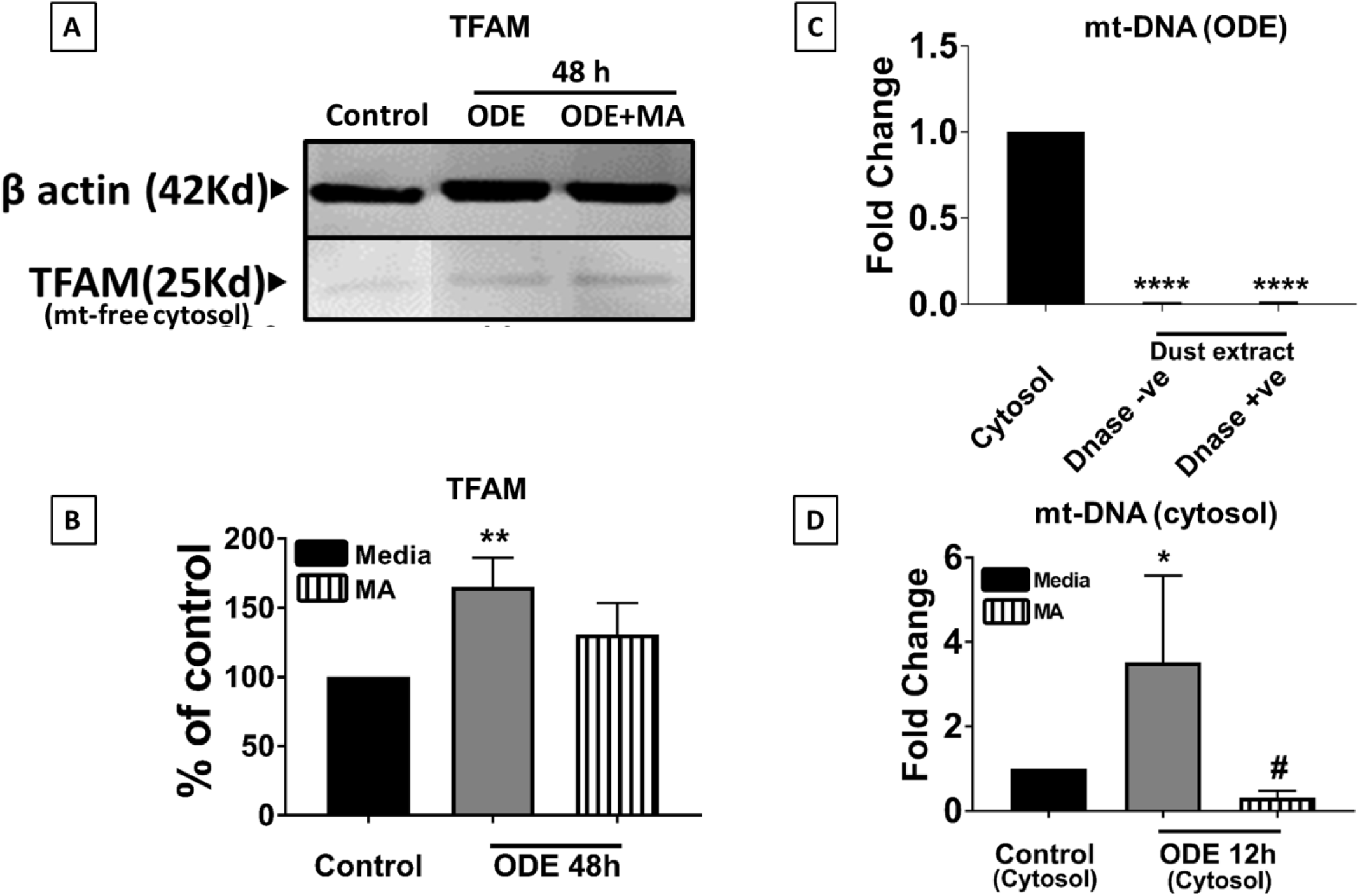
ODE induces the release of TFAM and mitochondrial DNA into the cytosol of microglia. Cells were exposed to media or ODE with or without MA and processed for Western blot analysis. Normalized (β-actin, loading control) bands of TFAM were processed for densitometry (ImageJ, NIH), and statistical analysis was performed. Compared to controls, ODE exposure significantly increased the TFAM levels at 48 h, and MA treatment had no effect (A-B). Mitochondria-free cellular cytosolic fraction and ODE treated with (negative control) or without DNase were processed for DNA extraction. mt-DNA specific primers were used for qRT-PCR analysis. ODE samples treated with and without DNAse confirmed that there was no background mitochondrial DNA in the ODE samples (C). Microglia treated with medium, ODE with or without mitoapocynin (C2) were processed to extract mitochondria-free cytosolic fraction, and mt-DNA content was quantified using qRT-PCR. Mitochondria free cytosol of ODE treated microglia contained significantly higher amounts of mt-DNA in the mitochondria free cytosolic fraction, and MA treatment significantly reduced the ODE induced mt-DNA release (D). (n=4, * exposure effect, # MA treatment effect and p < 0.05).

### ODE modulates TLR9 expression in microglia

Cultured microglial cells exposed to ODE with or without MA treatment were analyzed for TLR9 gene and protein expression. We employed qRT-PCR and found that in ODE treated microglia the gene expression of the TLR9 was highly upregulated as compared to control at 24 h. MA did not significantly change the TLR9 gene activity in ODE treated microglia (Fig. 7 A). To reinforce our findings, we also looked at the actual receptor protein expression in microglia using western blot analysis, and it followed a similar trend of upregulation upon ODE treatment (6 h, 24 h and 48 h) as compared to control. Surprisingly, in this case, MA was able to reduce the protein expression in contrast to the mRNA levels (MA data only for 48 h) (Fig. 7 B-C).

**Fig 7.**
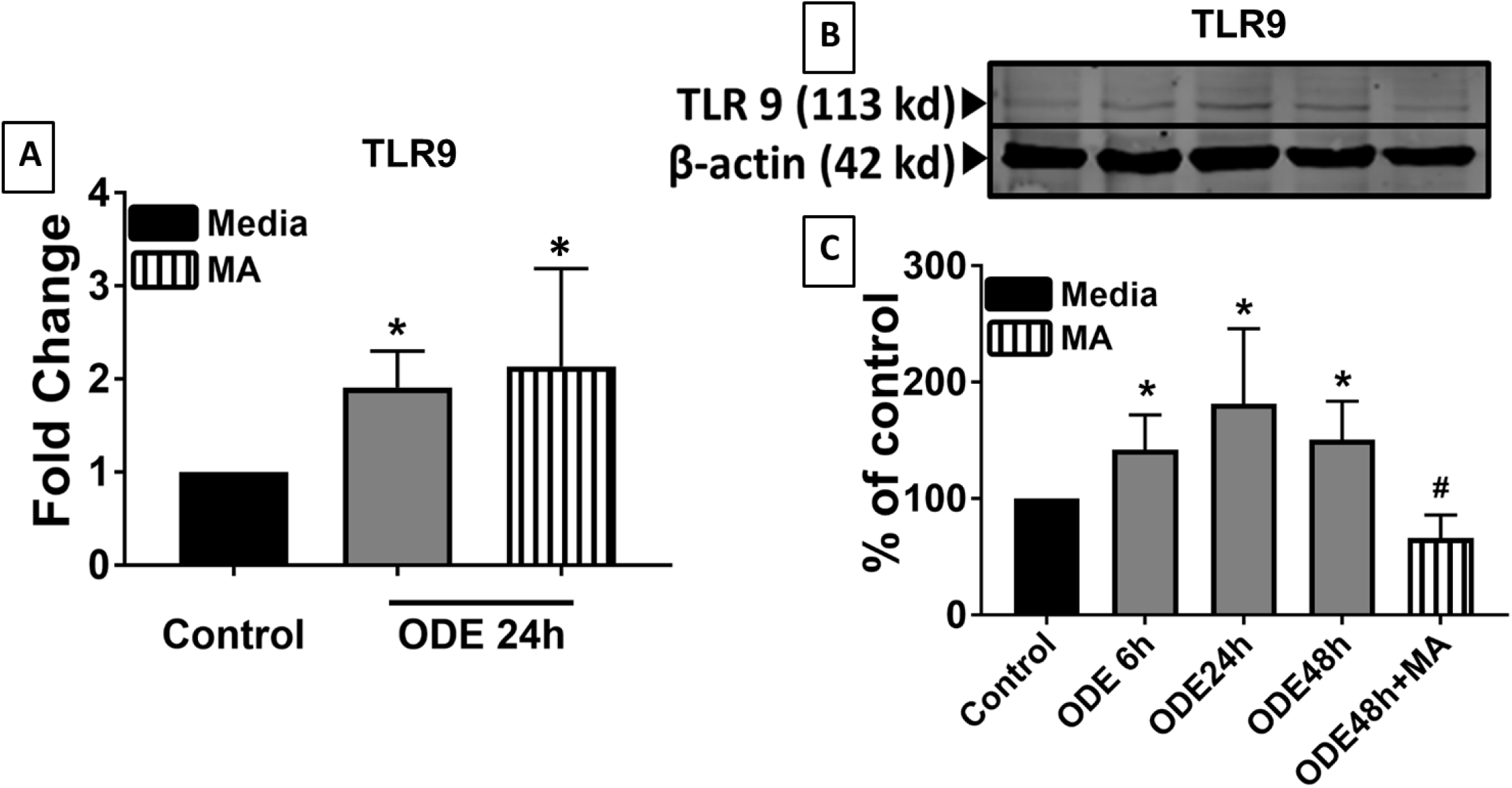
ODE exposure increases TLR9 expression. Microglia were exposed to media or ODE (6, 24, and 48 h) and co-treated with either vehicle or MA (48 h) and processed to quantify TLR9 mRNA (A) or protein levels (B-C). TLR9 specific primers were used to quantify mRNA (2^−ΔΔCt^ method). Compared to controls, TLR9 mRNA levels increased with ODE exposure (24 h), whereas MA treatment had no effect (A). TLR9 and β-actin antibodies (house-keeping protein) detected 113 and 42 kD bands, corresponding to TLR9 and β-actin respectively (B). Densitometry analysis of normalized bands showed that, compared to controls, ODE exposure increased the TLR9 levels at 6, 24, and 48 h, and MA treatment decreased the TLR9 levels at 48 h (C) (n=3, * exposure effect, # MA treatment effect, P < 0.05).

### ODE activates the cGAS-STING pathway

Cultured microglia exposed to ODE with or without MA treatment were processed for qRT-PCR and western blot for analyzing gene and protein expression respectively. Upon ODE treatment, we observed elevated mRNA levels of cGAS (12 h), STING (24 h), IRF3 (24 h), and IFN-β (24 h). MA was only able to reduce the mRNA levels of cGAS and STING following ODE treatment (Fig. 8 A-D). Protein levels of two end molecules of the pathway (cGAS and IFN-β) and were analyzed. Both cGAS and IFN-β protein levels were also found to be upregulated at 24 h and 48 h after ODE treatment, and MA did not affect cGAS and IFN-β protein levels at any given time point (Fig. 8 E-H).

**Fig 8.**
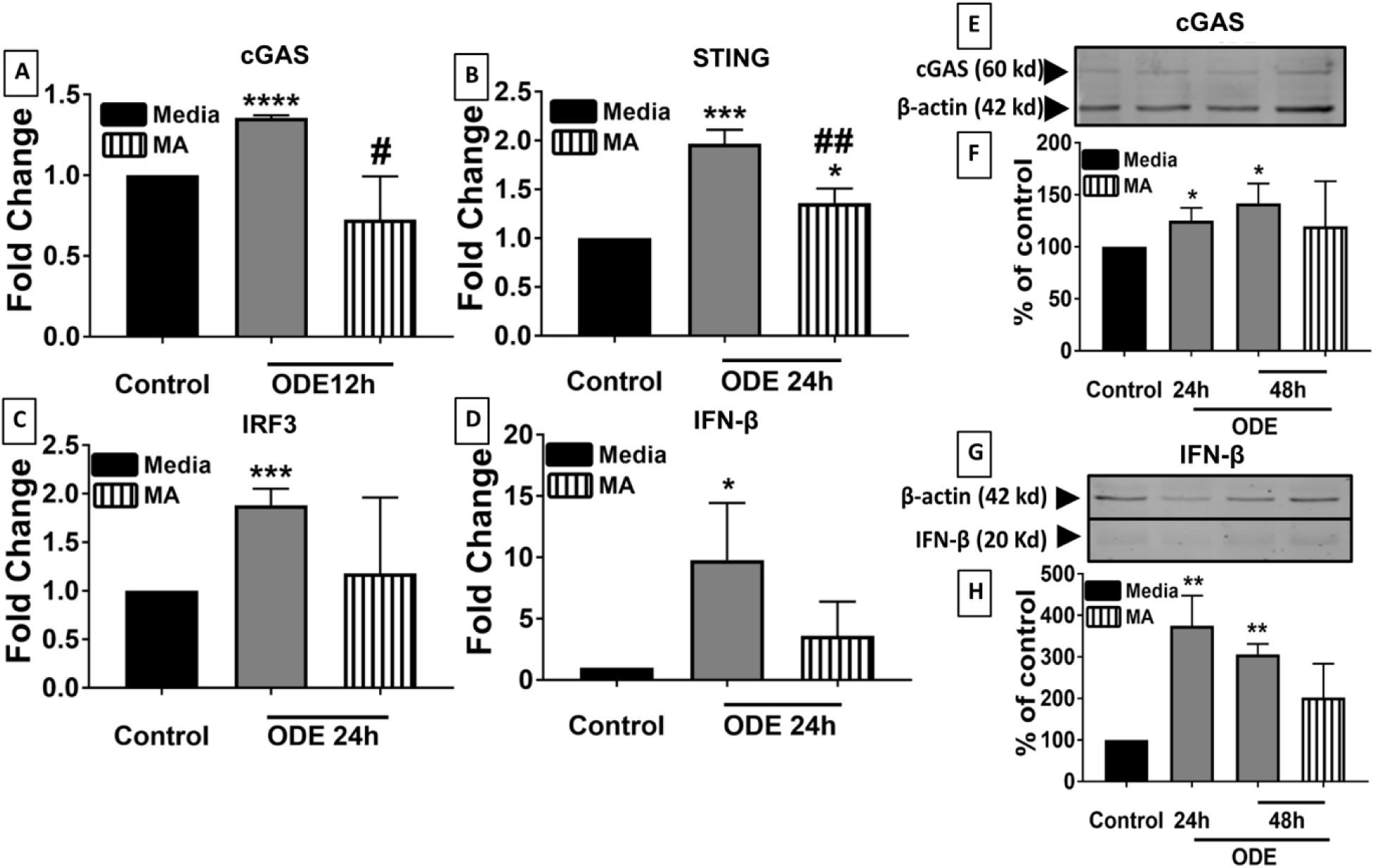
ODE exposure upregulates cGAS, STING, IRF3, and IFN-β expression in microglia. Microglia were exposed to media or ODE (12 h, 24 h, and 48 h) and co-treated with either vehicle or MA (48 h) and processed to quantify mRNA (A-D) or protein levels (E-H). cGAS, STING, IRF3, and IFN-β specific primers were used to quantify mRNA (2^−ΔΔCt^ method) and compared to controls, cGAS (12 h), STING (24 h), IRF3 (24 h) and IFN-β (24 h) mRNA levels increased. In contrast, MA treatment significantly decreased only cGAS and STING expression and did not affect IRF3 (C) and IFN-β (D) mRNA levels. cGAS (E), IFN-β (G) and β-actin antibodies (house-keeping protein) detected 60 kD, 20 kD, and 42 kD bands, respectively. Densitometry of normalized bands showed that, compared to controls, ODE exposure increased the cGAS(F) and IFN-β (H) levels at 24 and 48 h, and MA treatment had no effect (n=3, * exposure effect, # MA treatment effect, P < 0.05).

### cGAS-STING role in microglia activation

Successful transfection into microglia was confirmed by using a fluorescently tagged DsiRNA (DsiRNA TYE 563) at 10 nmol concentration after 24 h incubation (Fig. 9 A). After establishing effective transfection, microglia were incubated with R1, R2, or R3 at 10 nmol for 24 h concentration and then again exposed to either media (control) or ODE for further 48 h. A negative control group (NC) was also added using a scrambled DsiRNA at 10 nmol for 24 h. R1, R2, and R3 were successfully able to knockdown STING mRNA in ODE exposed microglia to the level of the control group (Fig. 9 B). Furthermore, IRF3 and IFN-β mRNA levels were also downregulated in ODE treated microglia to the level of control following treatment with R1, R2, or R3 (Fig. 9 C-D). After confirming an effective knockdown of STING and downregulation of IRF3 and IFN-β, we analyzed the expression of IBA-1 protein (microglial activation marker) in microglia. Our result indicated that R2 and R3 significantly reduced IBA-1 protein expression in microglia after ODE treatment (Fig. 9 E-F).

**Fig 9.**
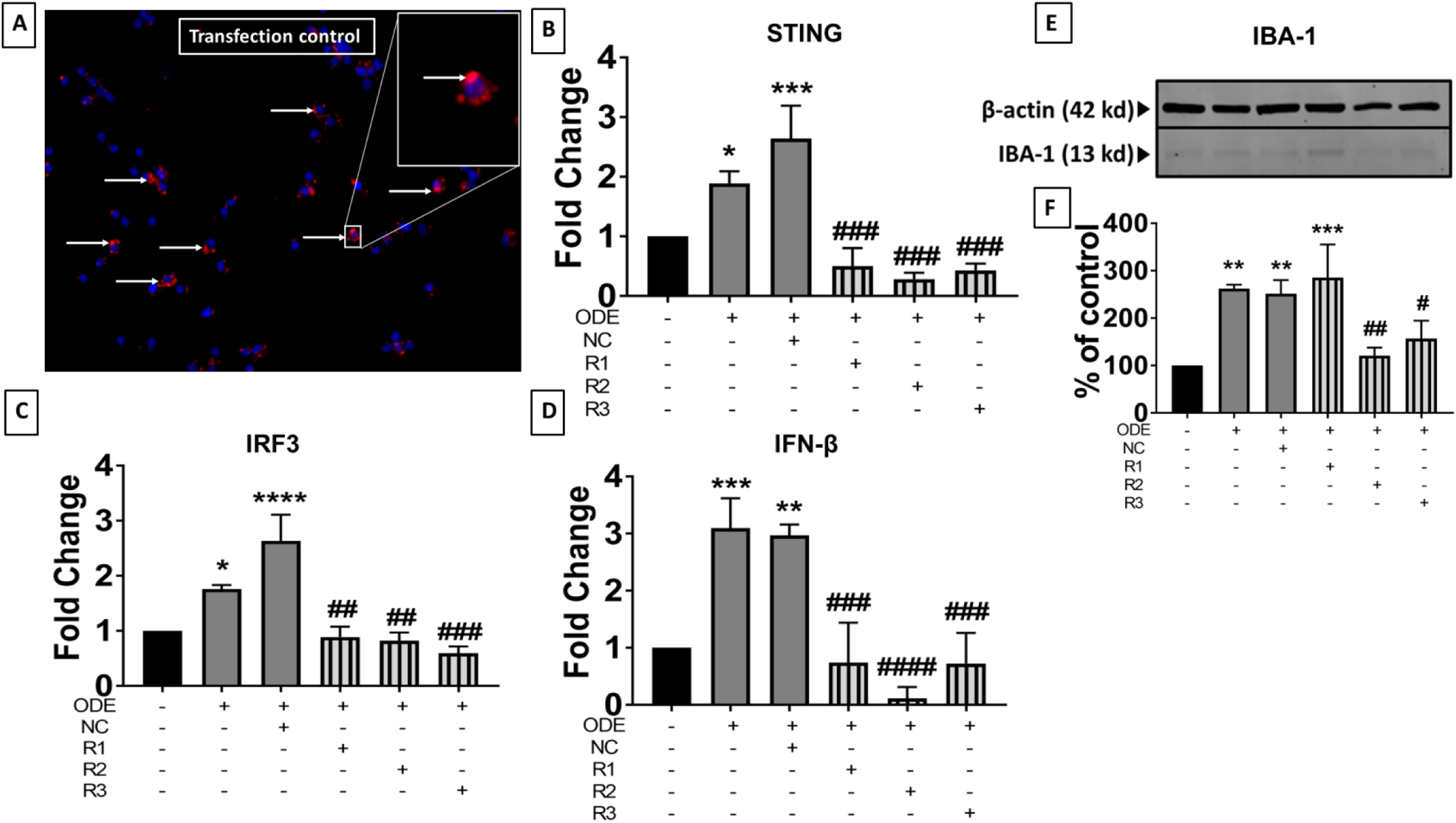
STING knockout with siRNA downregulates STING, IRF3, IFN-β, and IBA1 expression. Microglia were treated with DsiRNAs TYE 563 (transfection control, Cy3) or anti-STING siRNAs (R1, R2, R3) A scrambled siRNA was also used as a negative control (NC). Following treatment, cells were either fixed with paraformaldehyde (A) or processed for qRT-PCR analysis (B-D) or processed for western blot analysis (E-F). After 24 h, immunofluorescence (cy3, red) shows a successful transfection (white arrows and inset) (A) in the cytoplasm of microglia, and the nucleus is stained with DAPI (blue). Following qRT-PCR analysis, R1 (10 nmol), R2 (10 nmol), and R3 (10 nmol) significantly reduced the mRNA expression of STING (B), IRF-3 (C), and IFN-β (D) mRNAs at 24 h. Following the siRNA-mediated knockdown of STING mRNA, IBA-1 and β-actin (house-keeping protein) were detected in ODE treated (24 h) microglia at 13 kD and 42 kD bands, respectively (E). Normalized densitometry values show that, compared to ODE treated cells either with or without negative control siRNA (NC), anti-STING siRNA treatment (R2 and R3) reduced the IBA 1 protein levels at 24 h (F). (n=4, * exposure effect, # siRNA treatment effect, P < 0.05).

### ODE induces microglial activation in brain slice cultures (BSCs)

BSCs were prepared from mice pup brain and were cultured in the lab and treated with media (control) or ODE with or without MA(C11) (refer table 1) for 5 days. Compared to controls, Immunofluorescent staining showed high IBA-1 and reduced NeuN expression in the olfactory bulb, frontal cortex, corpus callosum, and visual cortex in ODE treated BSCs. MA(C11) treatment seems to have no effect on either IBA-1 and NeuN staining (Fig. 10 A). Following qRT-PCR analysis, expression of pro-inflammatory cytokines like TNF-α and IL-6 were found to be elevated with a significant rise in their gene activity after 5-day ODE treatment of BSCs when compared to control. MA(C11) significantly reduced both TNF-α and IL-6 expression in ODE treated BSCs (Fig. 10 B)

**Fig 10.**
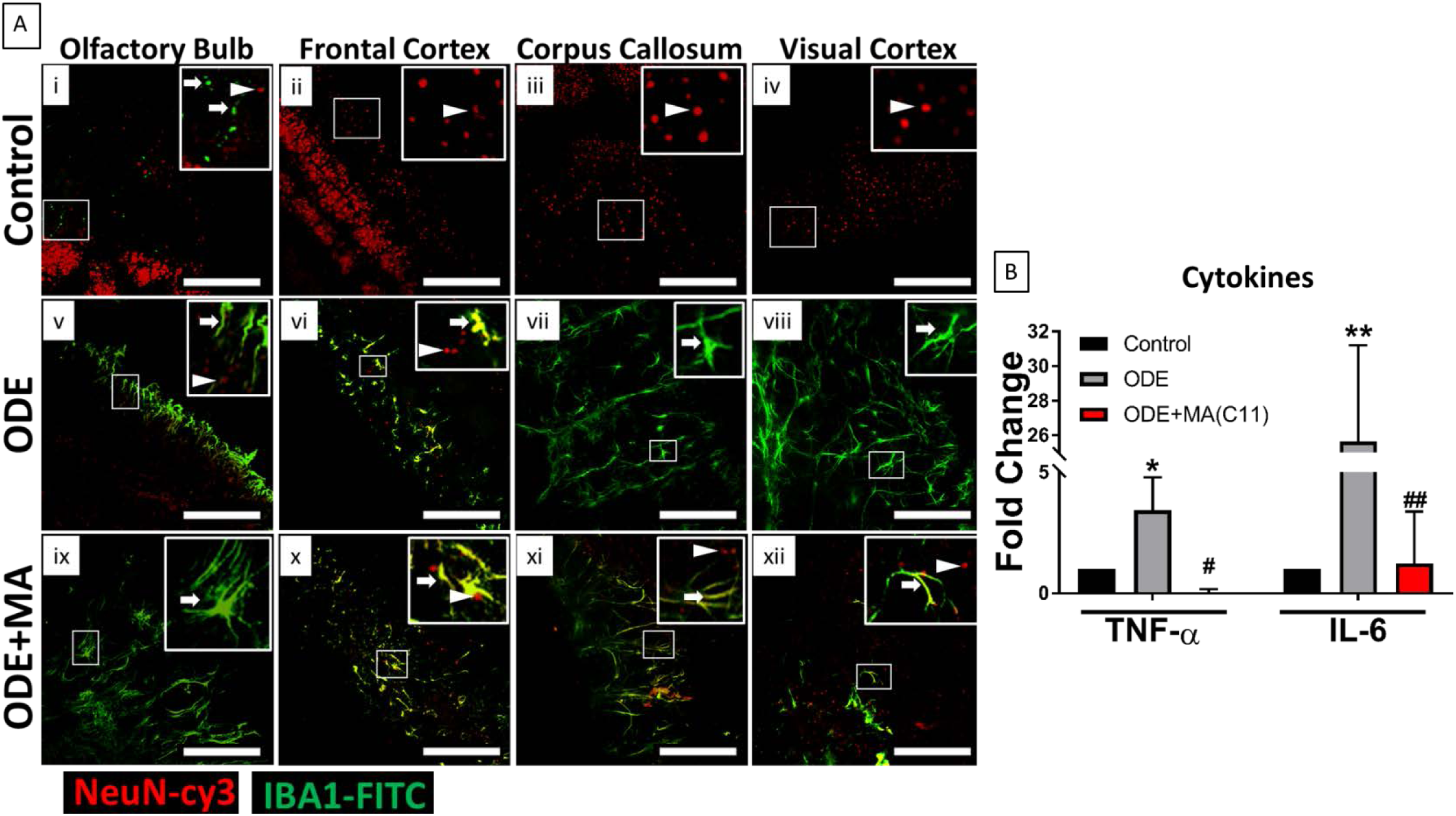
Organic dust extract (ODE)-exposure induces microglial activation and pro-inflammatory cytokines gene expression in organotypic brain slice culture (BSCs). BSCs were exposed to media (control) or ODE (5 days) followed by either vehicle or MA(C11) treatment and were stained with anti-NeuN (arrow head; neuronal marker; Cy3, red), anti-Iba1 (arrow; microglial activation marker; FITC, green) antibodies (A, i-xii) or processed for qRT-PCR analysis (B). Compared to control (A, i-iv), ODE-exposed BSCs showed higher amounts of Iba-1 staining in the olfactory bulb, frontal cortex, corpus callosum, and visual cortex of the brain (A, v-viii). MA(C11) treatment had no effect on ODE induced microglial activation (A, ix-xii). Compared to medium, ODE-exposed BSCs showed an upregulation of TNF-α and IL-6 gene expression. MA(C11) treatment decreased the gene expression of TNF-α and IL-6 (B). (n=3, * exposure effect, # MA(C11) treatment effect, P < 0.05, micrometer bar = 100 µm).

### ODE induces apoptosis in BSCs

TUNEL staining for dead or degenerating neurons was done in BSCs treated with media (control) or ODE with or without MA(C11) for 5 days. Fluorescent staining revealed that compared to control, ODE treated BSCs had a significantly higher number of degenerating or dead neurons in the olfactory bulb, frontal cortex and corpus callosum. Even BSCs co-treated with ODE and MA(C11) showed higher amounts of TUNEL positive cells (Fig. 11 A). The number of TUNEL positive cells were manually counted using Image J, and results were plotted (Fig. 11 B).

**Fig 11.**
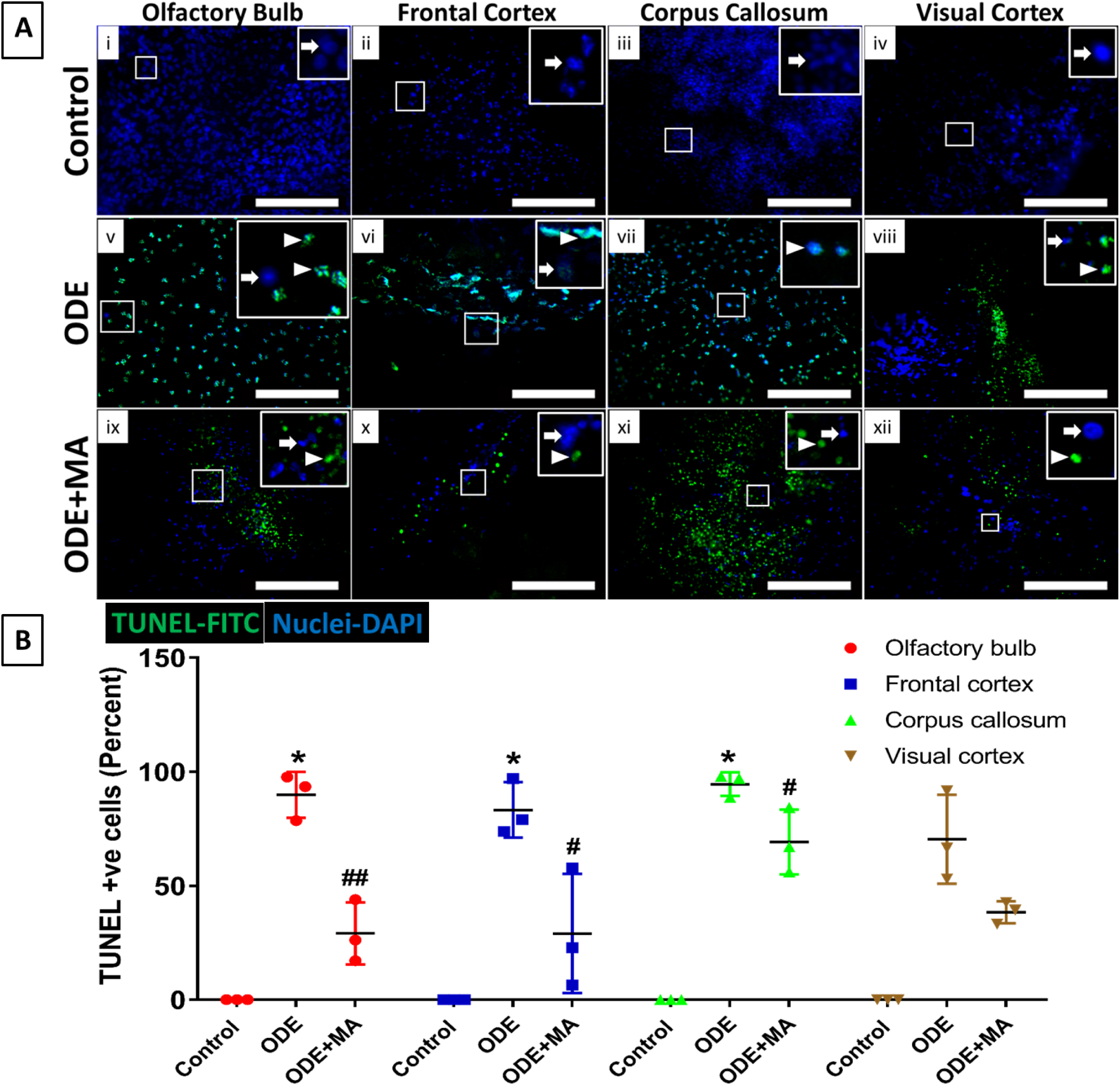
Organic dust extract (ODE)-exposure induces neurodegeneration in BSCs. BSCs were exposed to media (control) or ODE (5 days) and co-treated with either vehicle or MA (C11). BSCs were labeled with dUTP-FITC (arrow head; apoptosis marker, FITC, green), and nucleus stained with DAPI (arrow; blue) (A, i-xii). The total number of cells (DAPI, blue) and TUNEL positive cells (FITC, green) per field (20X) were counted in a total of five random fields. Compared to control (B, i-iv), ODE-exposed BSCs showed a higher number of TUNEL positive cells in the olfactory bulb, frontal cortex and corpus callosum regions of the brain (B, v-viii). MA(C11) significantly reduced the number of TUNEL positive cells in the olfactory bulb, frontal cortex, and corpus callosum. (B, ix-xii). (n=3, * exposure effect, # MA(C11) treatment effect, P < 0.05, micrometer bar = 100 µm).

### ODE induces mitochondrial damage in BSCs

Following 5-day treatment of BSCs with either media alone (control) or ODE with or without MA(C11), gene expression of MFN1, MFN2, DRP1, and PINK1 was analyzed. Compared to control, only MFN1 gene expression was found to be elevated in BSCs following 5 day ODE treatment. MA(C11) significantly reduced MFN1 gene expression in ODE treated BSCs (Fig. 12 A). Also, mt-DNA levels were found to be elevated in the cytosol of cells extracted from the ODE treated BSCs as compared to control BSCs. MA(C11) was again able to significantly prevent the release of mt-DNA in the cytosol (Fig. 12 B). However, no significant traces of mt-DNA were found in the culture media of both control or ODE treated BSCs (Fig.12 C).

**Figure 12.**
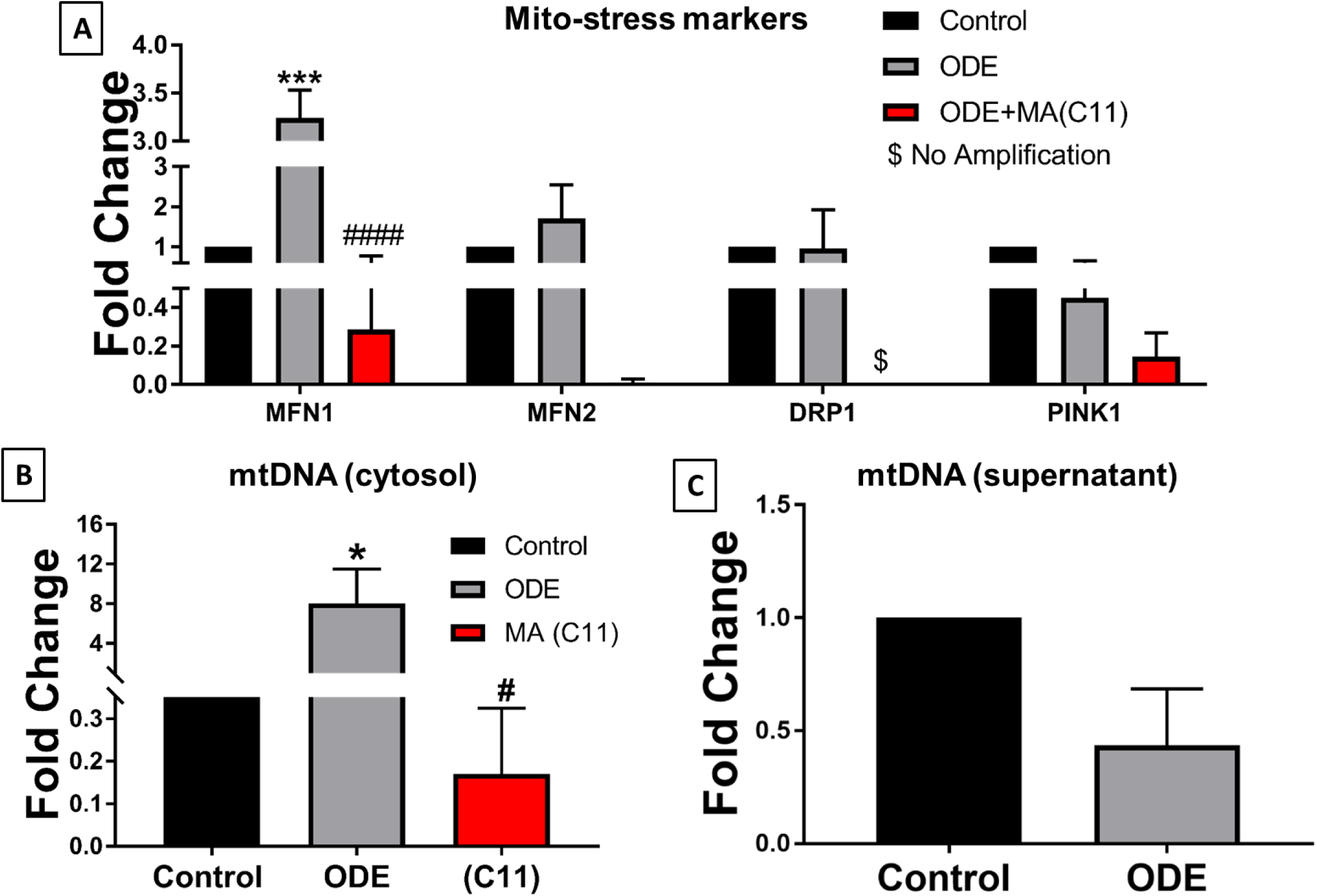
ODE exposure of organotypic brain slice culture activates mitochondrial stress response and induces the release of mt-DNA into the cytosol. BSCs were exposed to media or ODE (5 days) and co-treated with either vehicle or MA(C11). DNA extracted from the whole cell (A), mt-free cell cytosol (B), and supernatant (C) was processed for mt-DNA specific qRT-PCR analysis. Compared to medium, ODE-exposed brain slices showed an upregulation of MFN1 but not MFN2, DRP1, and PINK1. MA(C11) significantly decreased the MFN1 expression (A). Compared to control (vehicle), ODE induced a significant increase in the cytosolic mt-DNA fraction in BSCs. MA(C11) treatment significantly reduced cytosolic mt-DNA release in the cytosol (B). ODE exposed BSCs did not show any significant rise in mitochondrial DNA in supernatant (secreted) at 5 days post-treatment (C) (n=3, * exposure effect, # MA(C11) treatment effect, P < 0.05).

**Figure 13.**
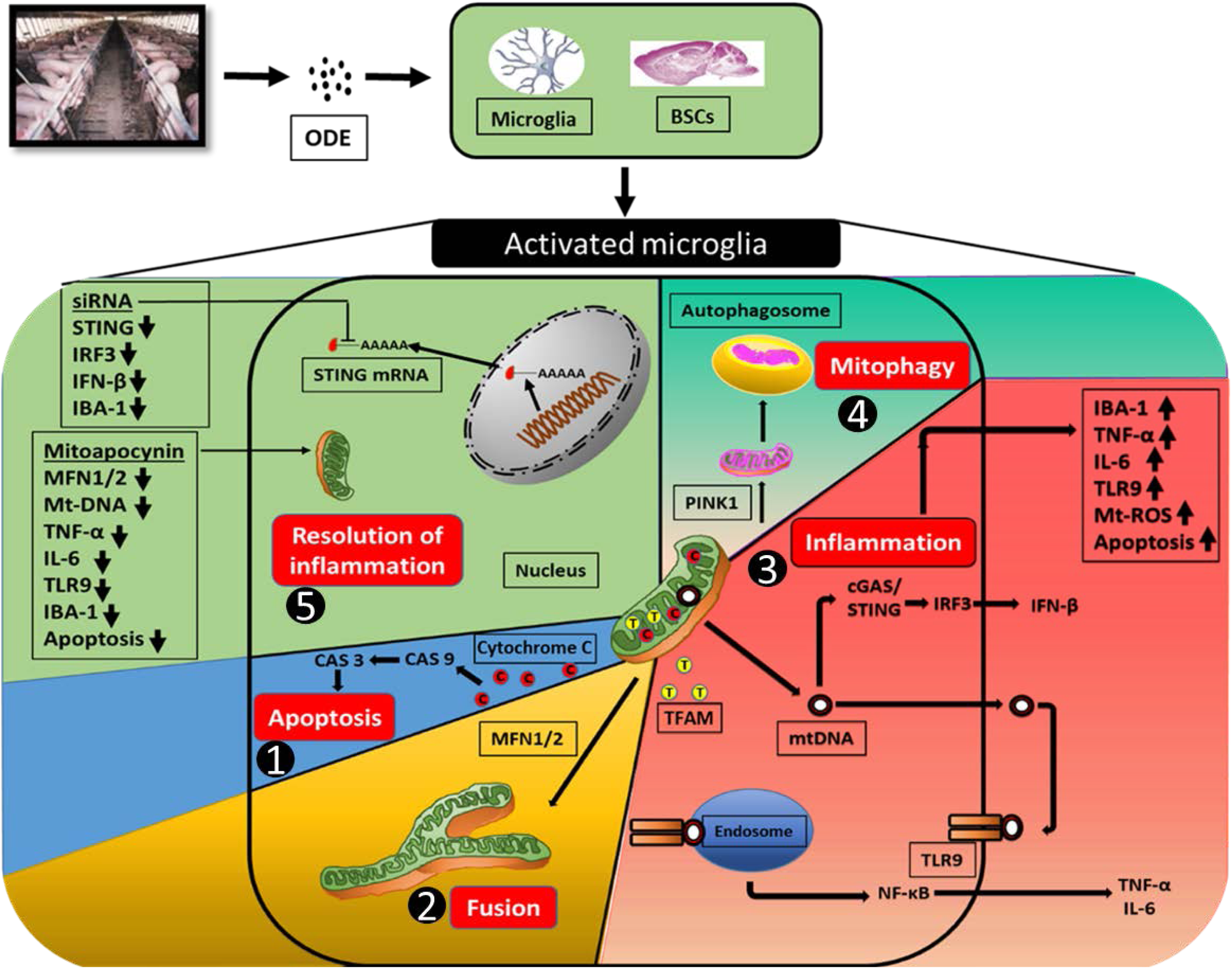
An overview of ODE exposure induced mitochondrial dysfunction in microglia and BSCs. Mitochondria is a vital organelle of the cell involved in the maintenance and survival of the cell. Thus, ODE induced mitochondrial damage can have different consequences simultaneously. 1) Cytochrome c is a respiratory chain protein loosely associated with the inner membrane of mitochondria. ODE exposure induced mitochondrial damage will result in the release of Cytochrome c into the cellular cytosol and initiation of apoptotic changes inside the cell through activation of Caspase 3 and Caspase 9. 2) Mitochondrial fusion (MFN1/2 mediated) is a form of stress response and is a mechanism for coping with the altered cellular homeostasis. 3) TFAM is a mitochondrial DNA binding protein that aids in the transcription of the mitochondria genome. Damage to mitochondria renders TFAM and mt-DNA vulnerable for release into cellular cytosol. When in the cytosol, mt-DNA can potentiate an inflammation response through TLR9-NFκB signaling resulting in pro-inflammatory cytokine release or mt-DNA can be sensed by cGAS-STING pathway, ultimately leading to IFN-β production. 4) PINK1 mediated mitophagy is a response often seen in damaged or stressed mitochondria to contain the inflammation. During mitophagy, mitochondria undergo selective degradation to keep the cell healthy. 5) Finally, by targeting mitochondria and preventing it from experiencing damage or stress can help alleviate the inflammation. MA (C2 or C11) helped in reducing mitochondrial fusion, preventing mt-DNA release, downregulating pro-inflammatory cytokines, preventing microglial activation and cellular apoptosis. Selective STING DsiRNA knockdown also helped in reducing microglial activation. Both MA (C2 or C211) and STING knockdown downregulated the ODE induced inflammatory and apoptotic markers and promoted the resolution of inflammation.

## DISCUSSION

Increasing incidences of neurocognitive disorders such as Parkinson’s disease among agricultural communities in the Midwestern and northeastern parts of the USA has been reported recently (Wright Willis *et al*., 2010b). The fact that the same area is known for higher density of animal production facilities is indicating a causal link between the agriculture exposures such as OD and incidences of Parkinson’s disease. We recently published a study and showed that organic dust extract (ODE) exposure activates microglia of the brain, leading to neuroinflammation and supported the hypothesis that OD if inhaled could induce neuroinflammation. Our data supported that mitochondria were involved in RNS production. Following treatment with MA (mitochondria targeted NOX-2 inhibitor), we were able to reduce inflammatory signals, and MA exposure also reduced microglial activation (Massey *et al*., 2019a). These evidences formed the basis for the current work that examined the impact of OD exposure on mitochondrial structure and function. In this manuscript, we investigated the role of mitochondria in the underlying mechanisms of ODE induced microglial activation and resultant neuroinflammation.

The OD exposure induced structural changes in the microglia (large amoeboid appearance and thicker branching) indicated their activation. Next, ultrastructural alterations in mitochondria indicated crystolysis, mitochondrial swelling, fusion, and increased size of structures resembling calcium sequestering bodies. We also noticed fragmentation and swelling of ER, increased expression of ER stress markers such as spliced XB1, ATF4 and GRP4 in microglia after ODE treatment. These findings attested that ODE treatment was inducing mitochondrial and ER stress responses in microglia. Interestingly, ODE treatment also induced mitochondrial bioenergetics deficiencies in agreement with the ultrastructural damages to the mitochondria. OD-induced upregulation of mitochondrial fusion genes (MFN1/MFN2) and PINK1 (apoptosis marker) indicated mitochondrial stress in ODE treated groups. Use of MA abrogated many of OD-induced damages at the gross, ultrastructural, bioenergetics and ER level indicating that OD-induced mitochondrial and ER damages could be abrogated by using MA.

The OD exposure induced cellular stress was marked an increase in mt-ROS and SOD-2 production. Further the mitochondrial oxidative stress was accompanied by an increase in the calcium uptake by mitochondria. Increased calcium update is known to altered mitochondrial membrane permeability that could lead to apoptotic changes in the cells. MA treatment did not reduce OD induced mt-ROS production, mitochondrial permeability and mitochondrial calcium uptake indicating that these changes are independent of mitochondria targeted NOX-2 inhibition (Langley, et al., 2017).

OD-exposure induced (TUNEL-staining positive) apoptotic cells to indicate the response to exposure could be protective since the apoptosis is a form of cell death that aims to contain the cell contents (such as DAMPs) in order to keep the inflammation in check (Kolb *et al*., 2017). We detected cytochrome c in the cellular cytosol (outside the mitochondria) along with an increase in caspase 3 and caspase 9 indicating that prevention of cytochrome c would be a good target (Oliva *et al*., 2016) to prevent exposure induced apoptosis. Surprisingly, MA treatment reduced the OD-induced increase in caspase-3 and 9 without an effect of the release of cytochrome c from the mitochondria. These results indicate that MA may have multiple anti-inflammatory effects.

Mitochondrial DNA resides within the mitochondrial matrix and is being identified as foreign or danger signal by the host immune system. Despite coding for very few mitochondrial genes, mt-DNA is important for oxidative phosphorylation (OXPHOS) and loss of mt-DNA is linked to several neurodegenerative diseases(Area-Gomez *et al*., 2019). Before examining if OD-exposure would lead to mt-DNA release or damage to mt-DNA, we verified that ODE samples used in our studies did not contain any DNA by using DNase treatment. ODE exposure of microglia resulted in the release of mt-DNA into the mt-free cellular cytosolic fraction and MA treatment significantly reduced the release of mt-DNA. ODE exposure increased the expression of TLR9, cGAS, STING as well as downstream molecules IRF-3 and IFN-β indicating that mt-DNA in the cytosol initiates a specific host response similar to a foreign DNA encounter. MA treatment reduced the expression of cGAS and STING but not the TLR9, IRF3 and IFN-β levels. mt-DNA is prokaryotic, perceived as foreign to the body and hence the immune system responds to the mt-DNA in a manner similar to foreign DNA through several intracellular sensing receptors such as TLR9 and cGAS molecules. Toll-like receptor 9 has a specific affinity for unmethylated cytosine and guanine nucleotides separated by a phosphate-backbone (CpG), which are common to prokaryotic DNA. This specificity is essential for preventing TLR9-dependent autoimmunity. mt-DNA also contains unmethylated CpG dinucleotides. TLR9 signaling can ultimately result in the activation of pro-inflammatory factors like NF-κB and cytokine secretion (McCarthy *et al*., 2015).

cGAS is a cytosolic sensor of foreign DNA that activates STING leading to IFN-β production. Activated cGAS produces cGAMP, which binds to STING, which resides in the ER. STING can further relay signals downstream to IRF3, ultimately leading to IFN-β production. Upon ODE exposure of microglia, we observed significant upregulation in the mRNA and protein molecules of the cGAS-STING pathway. MA again significantly reduced the upregulation of the cGAS and STING mRNAs. When we performed a specific knockdown of STING by using DsiRNA, it significantly reduced the IRF3 and IFN-β production as well as microglial activation as indicated by a reduced expression of IBA1. We found that MA treatment has produced promising results in curtailing mt-DNA release and its downstream signaling via TLR9 and cGAS-STING. This confirmed our hypothesis that mitochondria could serve as a crucial target to reduce OD-induced neuroinflammatory responses in microglia and mt-DNA release is a significant event as it compromises OXPHOS reactions and feeds into OD-induced inflammation.

In order to predict OD-exposure induced neuroinflammatory changes in the brain, we used mouse organotypic BSCs as a physiologically relevant model. ODE exposure of BSCs leads to the activation of microglia and the production of proinflammatory mediators such as TNF-α and IL-6. MA (C11) but not MA(C2) was effective in abrogating the ODE-induced expression of TNF-α and IL-6. These results suggested that MA(C11) that has been shown effective in *in-vivo* models is far superior when compared to MA(C2) which is suitable for *in vitro* models.

Similar to mouse microglia, ODE exposure of BSCs upregulated the IBA1 expression, pro-inflammatory cytokine production and induced TUNEL positive cells in different regions of BSCs. Further, ODE exposure of BSCs upregulated the expression of MFN1 (mitochondrial fusion protein), lead to release of mt-DNA into cytosol but not into the supernatant. Interestingly, MA(C11) was also able to reduce the ODE-exposure induced pro-inflammatory cytokine expression, MFN1 expression, and mt-DNA levels in cytosol to indicate that ODE-exposure induced mitochondrial dysfunction is a possible therapeutic target to treat neurodegenerative changes. In our study, we did not address the role of NLRP3 in mt-DNA induced signaling as well as the impact of loss of cells (TUNEL-positive) in certain regions of the brain (BSC model). The future studies addressing the behavioral, motor and sensory impacts of OD-exposure in animal models would be highly valuable.

In conclusion, OD-exposure through respiratory route activates microglial cells of the brain, induces mitochondrial and ER stress and inflammation characterized by the loss of cells in the brain. Exposure leads to mitochondrial dysfunction as indicated by expression of specific stress markers, structural and functional deficits and mt-DNA release and signaling. Use of mitochondria targeted NOX-2 inhibition (MA) or siRNA mediated knockdown of STING suppresses mt-DNA induced signaling. Both MA and STING appear promising in preventing OD-exposure induced neuroinflammation.

## CONCLUSIONS

Our results demonstrate that ODE-exposure, induces mitochondrial dysfunction and ultimate release of mt-DNA. Released mt-DNA is further triggers cGAs-STING pathway and leads to IFN-β production. Targeting cGAS-STING pathway or mitochondrial NOX-2 inhibition in ODE-induced microglial activation has the potential to reduce neuroinflammation and promote resolution.

## References

American Thoracic Society (1998). Respiratory health hazards in agriculture. In Am. J. Respir. Crit. Care Med. (Vol. 158, pp. S1–S76.

Anantharam, V., Kaul, S., Song, C., Kanthasamy, A., and Kanthasamy, A. G. (2007). Pharmacological inhibition of neuronal NADPH oxidase protects against 1-methyl-4-phenylpyridinium (MPP+)-induced oxidative stress and apoptosis in mesencephalic dopaminergic neuronal cells. Neurotoxicology 28(5), 988–997.

Area-Gomez, E., Guardia-Laguarta, C., Schon, E. A., and Przedborski, S. (2019). Mitochondria, OxPhos, and neurodegeneration: cells are not just running out of gas. The Journal of Clinical Investigation 129(1), 34–45.

Bhat, S. M., Massey, N., Karriker, L. A., Singh, B., and Charavaryamath, C. (2019). Ethyl pyruvate reduces organic dust-induced airway inflammation by targeting HMGB1-RAGE signaling. Respir. Res. 20(1), 27.

Block, M. L., Zecca, L., and Hong, J. S. (2007). Microglia-mediated neurotoxicity: uncovering the molecular mechanisms. Nat. Rev. Neurosci. 8.

Bronner, D. N., and O’Riordan, M. X. (2016). Measurement of Mitochondrial DNA Release in Response to ER Stress. Bio-protocol 6(12), e1839.

Chan, D. C. (2006). Mitochondria: Dynamic Organelles in Disease, Aging, and Development. Cell 125(7), 1241–1252.

Charavaryamath, C., and Singh, B. (2006). Pulmonary effects of exposure to pig barn air. J. Occup. Med. Toxicol. 1, 10.

Crews, F., and Vetreno, R. (2015). Mechanisms of neuroimmune gene induction in alcoholism. Psychopharmacology 233.

Cristóvão, A. C., Choi, D.-H., Baltazar, G., Beal, M. F., and Kim, Y.-S. (2009). The role of NADPH oxidase 1-derived reactive oxygen species in paraquat-mediated dopaminergic cell death. Antioxid Redox Signal 11(9), 2105–2118.

Dranka, B. P., Benavides, G. A., Diers, A. R., Giordano, S., Zelickson, B. R., Reily, C., Zou, L., Chatham, J. C., Hill, B. G., Zhang, J., et al. (2011). Assessing bioenergetic function in response to oxidative stress by metabolic profiling. Free Radical Biol Med 51(9), 1621–1635.

Dranka, B. P., Gifford, A., McAllister, D., Zielonka, J., Joseph, J., O’Hara, C. L., Stucky, C. L., Kanthasamy, A. G., and Kalyanaraman, B. (2014). A novel mitochondrially-targeted apocynin derivative prevents hyposmia and loss of motor function in the leucine-rich repeat kinase 2 (LRRK2(R1441G)) transgenic mouse model of Parkinson’s disease. Neurosci Lett 583, 159–164.

Escames, G., López, L. C., García, J. A., García-Corzo, L., Ortiz, F., and Acuña-Castroviejo, D. (2012). Mitochondrial DNA and inflammatory diseases. Hum. Genet. 131(2), 161–173.

Gao, H.-M., Liu, B., and Hong, J.-S. (2003). Critical role for microglial NADPH oxidase in rotenone-induced degeneration of dopaminergic neurons. J Neurosci 23(15), 6181–6187.

Ghosh, A., Langley, M. R., Harischandra, D. S., Neal, M. L., Jin, H., Anantharam, V., Joseph, J., Brenza, T., Narasimhan, B., Kanthasamy, A., et al. (2016). Mitoapocynin Treatment Protects Against Neuroinflammation and Dopaminergic Neurodegeneration in a Preclinical Animal Model of Parkinson’s Disease. J Neuroimmune Pharmacol 11(2), 259–278.

Halle, A., Hornung, V., Petzold, G. C., Stewart, C. R., Monks, B. G., Reinheckel, T., Fitzgerald, K. A., Latz, E., Moore, K. J., and Golenbock, D. T. (2008). The NALP3 inflammasome is involved in the innate immune response to amyloid-[beta]. Nat. Immunol. 9(8), 857–865 (10.1038/ni.1636).

Hinwood, M., Morandini, J., Day, T. A., and Walker, F. R. (2012). Evidence that Microglia Mediate the Neurobiological Effects of Chronic Psychological Stress on the Medial Prefrontal Cortex. Cereb. Cortex 22(6), 1442–1454.

Iowa State University and University of Iowa (2002). IOWA CONCENTRATED ANIMAL FEEDING OPERATIONS AIR QUALITY STUDY. Final Report. Available at: http://library.state.or.us/repository/2012/201204101013082/appendix_L.pdf. Accessed 07/10/2018 2018.

Jin, H., Kanthasamy, A., Ghosh, A., Anantharam, V., Kalyanaraman, B., and Kanthasamy, A. G. (2014). Mitochondria-targeted antioxidants for treatment of Parkinson’s disease: preclinical and clinical outcomes. Biochim Biophys Acta 1842(8), 1282–1294.

Johri, A., and Beal, M. F. (2012). Mitochondrial dysfunction in neurodegenerative diseases. The Journal of pharmacology and experimental therapeutics 342(3), 619–630.

Knoell, D. L., Smith, D. A., Sapkota, M., Heires, A. J., Hanson, C. K., Smith, L. M., Poole, J. A., Wyatt, T. A., and Romberger, D. J. (2019). Insufficient zinc intake enhances lung inflammation in response to agricultural organic dust exposure. J Nutr Biochem 70, 56–64.

Kolb, J. P., Oguin, T. H., 3rd, Oberst, A., and Martinez, J. (2017). Programmed Cell Death and Inflammation: Winter Is Coming. Trends Immunol. 38(10), 705–718.

Kondru, N., Manne, S., Greenlee, J., West Greenlee, H., Anantharam, V., Halbur, P., Kanthasamy, A., and Kanthasamy, A. (2017). Integrated Organotypic Slice Cultures and RT-QuIC (OSCAR) Assay: Implications for Translational Discovery in Protein Misfolding Diseases. Sci. Rep. 7(1), 43155.

Langley, M., Ghosh, A., Charli, A., Sarkar, S., Ay, M., Luo, J., Zielonka, J., Brenza, T., Bennett, B., Jin, H., et al. (2017). Mito-Apocynin Prevents Mitochondrial Dysfunction, Microglial Activation, Oxidative Damage, and Progressive Neurodegeneration in MitoPark Transgenic Mice. Antioxid Redox Signal 27(14), 1048–1066.

Lin, M. T., and Beal, M. F. (2006). Mitochondrial dysfunction and oxidative stress in neurodegenerative diseases. Nature 443(7113), 787–795.

Maekawa, H., Inoue, T., Ouchi, H., Jao, T.-M., Inoue, R., Nishi, H., Fujii, R., Ishidate, F., Tanaka, T., Tanaka, Y., et al. (2019). Mitochondrial Damage Causes Inflammation via cGAS-STING Signaling in Acute Kidney Injury. Cell reports 29(5), 1261–1273.e6.

Marzec, M., Eletto, D., and Argon, Y. (2012). GRP94: An HSP90-like protein specialized for protein folding and quality control in the endoplasmic reticulum. Biochim. Biophys. Acta 1823(3), 774–787.

Massey, N., Puttachary, S., Bhat, S. M., Kanthasamy, A. G., and Charavaryamath, C. (2019a). HMGB1-RAGE Signaling Plays a Role in Organic Dust-Induced Microglial Activation and Neuroinflammation. Toxicol. Sci. 169(2), 579–592.

Massey, N., Puttachary, S., Mahadev-Bhat, S., Kanthasamy, A. G., and Charavaryamath, C. (2019b). HMGB1-RAGE Signaling Plays a Role in Organic Dust-Induced Microglial Activation and Neuroinflammation. Toxicol. Sci. doi: 10.1093/toxsci/kfz071.

McCarthy, C. G., Wenceslau, C. F., Goulopoulou, S., Ogbi, S., Baban, B., Sullivan, J. C., Matsumoto, T., and Webb, R. C. (2015). Circulating mitochondrial DNA and Toll-like receptor 9 are associated with vascular dysfunction in spontaneously hypertensive rats. Cardiovasc. Res. 107(1), 119–30.

Motwani, M., Pesiridis, S., and Fitzgerald, K. A. (2019). DNA sensing by the cGAS–STING pathway in health and disease. Nat Rev Genet 20(11), 657–674.

Nakayama, H., and Otsu, K. (2018). Mitochondrial DNA as an inflammatory mediator in cardiovascular diseases. The Biochemical journal 475(5), 839–852.

Nath Neerukonda, S., Mahadev-Bhat, S., Aylward, B., Johnson, C., Charavaryamath, C., and Arsenault, R. J. (2018). Kinome analyses of inflammatory responses to swine barn dust extract in human bronchial epithelial and monocyte cell lines. Innate Immun. 24(6), 366–381.

Nordgren, T. M., and Charavaryamath, C. (2018). Agriculture Occupational Exposures and Factors Affecting Health Effects. Curr. Allergy Asthma Rep. 18(12), 65.

Oliva, C. R., Markert, T., Ross, L. J., White, E. L., Rasmussen, L., Zhang, W., Everts, M., Moellering, D. R., Bailey, S. M., Suto, M. J., et al. (2016). Identification of Small Molecule Inhibitors of Human Cytochrome c Oxidase That Target Chemoresistant Glioma Cells. J Biol Chem 291(46), 24188–24199.

Oslowski, C. M., and Urano, F. (2011). Measuring ER stress and the unfolded protein response using mammalian tissue culture system. Methods Enzymol. 490, 71–92.

Poole, J. A., Thiele, G. M., Janike, K., Nelson, A. J., Duryee, M. J., Rentfro, K., England, B. R., Romberger, D. J., Carrington, J. M., Wang, D., et al. (2019). Combined Collagen-Induced Arthritis and Organic Dust-Induced Airway Inflammation to Model Inflammatory Lung Disease in Rheumatoid Arthritis. J. Bone Miner. Res. doi: 10.1002/jbmr.3745.

Ransohoff, R. M. (2016). How neuroinflammation contributes to neurodegeneration. Science 353(6301), 777–783.

Riley, J. S., Quarato, G., Cloix, C., Lopez, J., O’Prey, J., Pearson, M., Chapman, J., Sesaki, H., Carlin, L. M., Passos, J. F., et al. (2018). Mitochondrial inner membrane permeabilisation enables mtDNA release during apoptosis. The EMBO journal 37(17), e99238.

Roy, S. R., Schiltz, A. M., Marotta, A., Shen, Y., and Liu, A. H. (2003). Bacterial DNA in house and farm barn dust. J. Allergy Clin. Immunol. 112(3), 571–8.

Truban, D., Hou, X., Caulfield, T. R., Fiesel, F. C., and Springer, W. (2017). PINK1, Parkin, and Mitochondrial Quality Control: What can we Learn about Parkinson’s Disease Pathobiology? J. Parkinsons Dis. 7(1), 13–29.

Warren, K. J., Dickinson, J. D., Nelson, A. J., Wyatt, T. A., Romberger, D. J., and Poole, J. A. (2019). Ovalbumin-sensitized mice have altered airway inflammation to agriculture organic dust. Respir. Res. 20(1), 51.

Wolf, S. A., Boddeke, H. W., and Kettenmann, H. (2017). Microglia in Physiology and Disease. Annu Rev Physiol 79, 619–643.

Wright Willis, A., Evanoff, B. A., Lian, M., Criswell, S. R., and Racette, B. A. (2010a). Geographic and ethnic variation in Parkinson disease: a population-based study of US Medicare beneficiaries. Neuroepidemiology 34(3), 143–51.

Wright Willis, A., Evanoff, B. A., Lian, M., Criswell, S. R., and Racette, B. A. (2010b). Geographic and Ethnic Variation in Parkinson Disease: A Population-Based Study of US Medicare Beneficiaries. Neuroepidemiology 34(3), 143–151.

